# Drought and rewetting drive divergent microbial dynamics across soil compartments and reveal *Trinickia* as a key genus for soybean drought resilience

**DOI:** 10.64898/2026.09.15.751935

**Authors:** Timothy M. Ghaly, Vanessa J. McPherson, Vaheesan Rajabal, Elena Colombi, Sasha G. Tetu

## Abstract

Drought conditions are projected to intensify under climate change, and engineering more resilient plant microbiomes is a promising route to supporting crop productivity. However, effectively harnessing these microbiomes depends on identifying keystone taxa and functional traits that promote plant drought tolerance. Here, we combine deep shotgun metagenomics with absolute-abundance calibration to track genome-resolved microbial dynamics across a prolonged soybean drought and rewetting cycle. Leveraging 728 high-purity metagenome-assembled genomes that capture ∼83% of prokaryotic reads on average, we show that drought drives significantly greater change in community structure in the rhizosphere than in bulk soil. This was accompanied by declining taxonomic, yet rising functional diversity. While rhizosphere communities rapidly recovered within 24 hours of rewetting, bulk soil communities instead underwent a progressive post-rewetting disturbance, consistent with a Birch-effect-driven successional shift. Rhizosphere drought enrichment was dominated by a single, taxonomically underexplored genus, *Trinickia*, whose genomes encode a coordinated suite of osmotic-stress-tolerance, bio-fertilisation, and phytohormone/polyamine biosynthesis traits consistent with plant growth promotion. Supporting this genomic inference, *Trinickia* abundance strongly correlated with increased root biomass during active drought, independent of plant developmental stage. Together, these findings identify the rhizosphere as a dynamically responsive compartment during drought and rewetting, and highlight *Trinickia* species as genomically and phenotypically supported candidates for microbiome-based strategies to improve soybean drought resilience.

## Introduction

Drought is a major driver of global crop production loss, and its frequency and severity are projected to intensify under continued climate change [1]. Among staple crops, soybean is disproportionately vulnerable, with recent global crop modelling projecting its yield losses to exceed those of other major crops, with an average country-level reduction of 16.4% by 2050 [1]. As a globally significant legume crop underpinning both protein supply and biological nitrogen input to agricultural soils, understanding how drought reshapes its supporting root microbiome is urgently needed.

Root-associated microbes can buffer host plants against drought stress through a range of established mechanisms. These include the biosynthesis of phytohormones such as indole-3-acetic acid (IAA), which promotes root growth and architectural changes that improve water foraging [2–5]; the mobilisation of nutrients, including nitrogen, phosphorus, and iron, that become less bioavailable to plants under water deficit [6]; and the production of compatible solutes and polyamines, which help maintain cellular osmotic balance and limit oxidative damage under water stress in both the host plant and the microorganisms themselves [7, 8]. Collectively, these functions position the rhizosphere microbiome as an important resource for improving crop drought resilience [9].

Soil microbial communities are themselves fundamentally reshaped by water limitation, with cascading effects on the biogeochemical processes that they mediate [10–12]. Drought is well known to drive compositional shifts in rhizosphere microbiomes [13–15], with these effects greater in the rhizosphere than the surrounding soil [16–18]. A major driver of this pronounced rhizosphere restructuring is the plant’s own drought response: water deficit alters the composition of root exudates and the polysaccharide structure of root mucilage in ways that reshape substrate availability for the colonising microbial community [19–22]. This stress-induced metabolic shift is thought to selectively recruit root-associated microbiota that can enhance host drought tolerance, forming a positive host-microbiome feedback loop [22–24].

However, identifying which specific microbial taxa are recruited under drought conditions, and how their functional traits mediate host drought tolerance remains challenging. To date, taxonomic profiling via amplicon sequencing has helped resolve community composition but left the functional gene content of individual organisms uncharacterised [13, 15], while previous genome-resolved analyses have captured only a tiny fraction of the total community due to insufficient sequencing resolution [14]. Fully realising the functional potential of rhizosphere microbiomes requires approaches that go beyond tracking relative taxonomic shifts to resolving absolute population dynamics at high genomic resolution. Relative abundance metrics alone can obscure whether a proportional increase reflects true population expansion or merely the decline of co-occurring taxa [25]. Therefore, incorporating absolute-abundance calibrations provides crucial quantitative insights needed to trace true population trajectories through drought and recovery phases.

Here, we track these dynamics using a controlled glasshouse drought and rewetting experiment in soybean, applying deep shotgun metagenomic sequencing across paired rhizosphere and bulk soil compartments. By combining comprehensive genome recovery with absolute-abundance calibration, we capture microbial community and individual-genome trajectories through both stress and recovery phases. We use this framework to: (i) characterise shifts in taxonomic and functional diversity across soil compartments and drought phases; (*ii*) identify key microbial populations and functional traits driving these responses at genome resolution; and (*iii*) evaluate whether genomically inferred plant-growth-promoting traits translate into measurable host phenotypic outcomes.

## Methods

### Glasshouse soybean drought experiment

The plant growth experiment was conducted using Soybean (*Glycine max* cv. Hayman) grown in potted topsoil (Burgess Soil, Wallacia, NSW, Australia). Triplicate soil samples were taken prior to the experiment for physicochemical analysis conducted at the Environmental Analysis Laboratory (Southern Cross University, Lismore, Australia). The soil was characterised as near-neutral (pH 7.09), and possessed a low nutrient-retention capacity (ECEC: 6.35 cmol+/kg) alongside low baseline levels of total carbon (0.96%), nitrogen (0.07%), available phosphorus (19.4 mg/kg), and exchangeable potassium (71.4 mg/kg).

Unsterilised soybean seeds were sown at a depth of 3 cm in tubestock seedling trays containing the topsoil. Seedlings were maintained under an automated overhead sprinkler system configured to deliver a 1-minute watering duration three times daily at 07:00, 12:00, and 16:00. After 13 days of growth, the seedlings were transplanted into experimental pots (top dimensions: 17 × 17 cm, bottom dimensions: 14 × 14 cm, height: 23cm) containing the same topsoil. Environmental conditions within the glasshouse were set to approximately replicate the climate profile of the Burnett soybean growing region in southeastern Queensland (Kingaroy baseline; Bureau of Meteorology data; https://www.bom.gov.au/climate/averages/tables/cw_040922_All.shtml). The diurnal temperature cycle ranged from 18°C to 31°C (06:00–19:30), with the maximum temperature threshold maintained for a 4.5-hour block over midday. Nocturnal temperatures were held at a constant 18°C (19:30–06:00).

Following a 6-week plant establishment phase, a 4-week drought treatment (T1–T3) was imposed using a capillary irrigation system designed to simulate a controlled, continuous water deficit [26]. This approach involves placing potted plants on top of a column of porous foam within tubs maintained at a desired water level, where a greater vertical distance between the base of the pot and the water table produces a more intense drought (Supplementary Fig. S1). The method relies on capillary action to apply a gradual water deficit, rather than typical cyclic drying-and-rewetting profiles. Control pots were positioned on 23-cm columns of commercial porous foam (STRASS® IDEAL floral foam bricks) inside plastic tubs, with the water level strictly maintained at a height that was level with the top of the foam blocks to prevent soil water deficit (Supplementary Fig. S1). In contrast, drought-treated pots were maintained in separate tubs with the water table maintained at a distance of 10 cm from the top of the foam blocks to establish and sustain a constant water deficit (Supplementary Fig. S1). Baseline sampling (T0) was conducted one day prior to drought initiation. During the active drought phase, the automated overhead sprinkler system was deactivated, while the respective water depths within both control and drought tubs were continuously maintained. The drought phase spanned four weeks, with destructive harvests conducted at 7 days (T1), 19 days (T2), and 28 days of drought (T3), sampling six control and six treatment replicate pots per time point. Pots were randomly re-positioned twice weekly throughout the duration of the experiment.

To initiate the recovery phase, all remaining pots were removed from the tubs, and the automated overhead sprinkler regime was returned to match the establishment conditions. Recovery sampling was performed at 1 day (T4), 4 days (T5), and 14 days (T6) post-rewetting, using six replicates per treatment group at each time point.

### Sample collection and processing

For sampling, bulk and rhizosphere soil fractions of each plant were isolated concurrently. To sample bulk soil, the top ∼1 cm of surface material was cleared from a location approximately 1 cm away from the outer edge of the pot using an ethanol-sterilized metal scoop. A 3-cm core of soil directly beneath this cleared layer was collected into 5 mL tubes and stored at –20°C until DNA extraction.

To isolate the rhizosphere fraction, the root system of each plant was shaken vigorously to dislodge loose soil. The remaining root system, with closely-adhered rhizosphere soil, was placed inside sterile 50 mL tubes filled with phosphate-buffered saline containing 0.05% (v/v) Tween-20 (PBST). To dislodge the rhizosphere soil fraction, tubes were placed in a water bath sonicator and sonicated at 40 kHz for 10 minutes, with ice. This was followed by horizontal orbital shaking at 250 rpm for 2 minutes. The PBST rhizosphere soil suspension was centrifuged at 3,200 × *g* for 10 mins to pellet the soil fraction. Pellets were resuspended in 1.5mL of PBST and stored at −20°C for subsequent DNA extraction. Root nodules were observed but were not directly sampled.

To determine gravimetric soil moisture at each harvest point, ∼100g of homogenised soil from each pot was transferred to a foil tray immediately after plant harvesting and weighed to establish fresh weight. The soil samples were transferred to a drying oven at 80°C for a minimum of 2 days and then re-weighed. Gravimetric soil water content (GWC, %) was determined using the mass profiles of the soil samples according to the following formula: GWC = (Fresh soil weight –Dry soil weight) ÷ Dry soil weight. Above-ground shoot biomass and washed root systems were also dried at 80°C for a minimum of 2 days and weighed to obtain dry weights (Supplementary Table S1).

Rhizosphere and bulk soil DNA extractions were performed using a previously established bead-beating protocol [27]. Three of the six replicates per treatment group at each time point were randomly selected for whole metagenomic shotgun sequencing. This resulted in a total of 78 samples (39 rhizosphere and 39 bulk soil samples) sequenced on an Illumina NovaSeq X Plus 10B (300-cycle) platform at the Ramaciotti Centre for Genomics (UNSW, Sydney, Australia). Sequencing produced on average 109 million × 150-bp paired-end reads (16.3 Gbp) per sample. All sequence data have been made available via the European Nucleotide Archive (ENA) under study accession PRJEB102647 (BioSamples: SAMEA120513386 –SAMEA120513463).

### Sequence read processing

Paired-end metagenomic reads underwent adapter-clipping and quality-trimming using fastp v0.24.1 [28, 29], employing a 4bp sliding window, from 5’- to 3’-end, with a mean quality threshold of 20 [parameters: --cut_right --cut_right_window_size 4 --cut_right_mean_quality 20 --thread 16]. Quality-trimmed reads were then cleaned to remove any contaminating host (soybean), human, or Illumina PhiX control spike-in reads. This involved mapping the trimmed reads using Strobealign v0.17.0 [30] to the telomere-to-telomere genomes of soybean (*Glycine max* Williams 82 cultivar, GCA_030864155.1) [31] and human (T2T-CHM13+Y v2.0) [32, 33], as well as the *Escherichia* phage phiX174 genome (NC_001422.1). Paired-end reads where both read pairs mapped to a reference genome were identified with SAMtools [34], and removed using the *filterbyname* program from the BBTools v38.93 software suite (https://sourceforge.net/projects/bbmap/). The resulting high-quality, filtered reads were used for all downstream analyses, establishing a total dataset of 7.8 billion reads (1.2 terabases) with an average of 99.8 million reads (14.7 Gbp) per sample (Supplementary Table S2).

### Taxonomic and functional profiling of metagenomic reads

Taxonomic profiling of metagenomic reads was performed using SingleM v0.20.3 [35], which estimates community composition by identifying reads spanning conserved regions within universal single-copy marker genes (SingleM database v5.4.0). Taxonomy is then assigned based on the Genome Taxonomy Database (GTDB) release 226 [36–39]. The SingleM *pipe* subcommand was run with default parameters on each sample individually. SingleM *summarise* was then run to generate OTU tables, clustered at 96.67%, for each universal marker across all samples. OTU count data were then normalised by total-sum scaling (TSS). For downstream alpha- and beta-diversity analyses, we used all ribosomal protein marker tables (*n*=14 markers). Bray-Curtis dissimilarities [40] and Shannon diversity indices [41] were calculated separately for each of the 14 ribosomal marker tables using the *vegdist* and *diversity* functions, respectively, from the vegan R package v2.5.7 [42], both run with default parameters. The resulting values were averaged across all markers to obtain a single consensus per sample (Shannon diversity) or sample pair (Bray-Curtis distance). The average prokaryote genome size per sample was inferred using SingleM *prokaryotic_fraction* [43] with default settings, which calculates the total bases derived from bacteria and archaea using reference genome sizes from each taxon’s GTDB lineage. Taxonomic profiling of microeukaryotes was performed using EukDetect v2.0.1 [44] with default parameters.

Functional profiling of metagenomic reads was performed using HUMAnN v3.9 [45]. For this, paired-end reads were first concatenated into a single file for each sample, and profiled individually with HUMAnN 3 using default parameters. The resulting pathway abundance files were normalised to relative abundances using the *humann_renorm_table* script and merged using *humann_join_tables*. Finally, the *humann_split_stratified_table* script was used to separate stratified rows (taxon-specific contributions) from unstratified rows (community-level totals).

### Metagenomic assembly and genome binning

We employed a co-assembly approach, which has been shown to enhance assembly contiguity [46] and improve the recovery of metagenome-assembled genomes (MAGs) [47]. Given the massive size of the dataset (1.2 terabases), we performed two separate metagenomic co-assemblies, one for all treatment samples, and another for control samples. For each co-assembly, pooled forward and pooled reverse reads were depth-normalised using BBNorm v38.93 (https://sourceforge.net/projects/bbmap/) to a target *k*-mer depth of 70, and min depth of 2 [parameters: target=70 mindepth=2 prefilter=t threads=48]. Normalised reads were then co-assembled using MetaHipMer2 [46] with default parameters. MetaHipmer2 is uniquely capable of performing such massive co-assemblies by scaling efficiently on distributed-memory, multi-node supercomputers. We executed MetaHipMer2 across 250 HPC nodes on the Gadi supercomputer (National Computational Infrastructure Australia), using 12,000 CPU cores and 47.5 terabytes of memory. Each co-assembly was filtered to discard any contigs shorter than 1 kb.

We used the metagenomic co-assemblies to perform multi-sample binning using three different binning tools: COMEBin v1.0.4 [48], GenomeFace [49] and MetaBAT 2 v2.17 [50]. First, contig coverage was calculated using CoverM v0.7.0 [51] by mapping the filtered reads from each sample to the co-assembled contigs with Strobealign [parameters: contig -p strobealign --strobealign-use-index --min-read-percent-identity 97 -m metabat --bam-file-cache-directory]. For GenomeFace and MetaBAT 2, the CoverM contig coverage profiles were provided alongside the co-assemblies, using the default minimum contig length thresholds of 1,500 bp for GenomeFace, and 2,500 bp for MetaBAT2. For COMEBin, coverage was supplied via sample-specific BAM files generated by CoverM, using a filtered subset of contigs with a minimum length of 2,500 bp. Both COMEBin and GenomeFace were executed in GPU mode using four Nvidia Tesla Volta V100 GPUs and 48 CPU cores. The output bins from the three binning tools were then merged and refined using Binette v1.2.1 [52] with default settings, which generates optimised hybrid bins by identifying overlapping contig assignments across the different bin sets and retaining the combinations that maximise overall genome quality, assessed using CheckM2 v1.1.0 [53]. This binning workflow was run for both co-assemblies separately, and then pooled and dereplicated at 95% average nucleotide identity (ANI) using Galah [54], as part of the CoverM package. Galah was run using skani [55] for ANI calculations, and the CheckM2 output was used for genome quality information [parameters: coverm cluster --ani 95 --checkm2-quality-report --precluster-method skani --cluster-method skani]. Only high-purity MAGs, with CheckM2-estimated contamination <5% and completeness > 50%, were retained for downstream analysis. This workflow recovered 728 MAGs with a median completeness of 71.56% and a median contamination of 1.08% (Supplementary Table S3). MAGs were taxonomically classified with GTDB-Tk v2.4.1[56, 57] against GTDB release 226 [parameters: classify_wf --skip_ani_screen].

### Absolute abundance calibration of metagenome-assembled genomes

To overcome the inherent limitations of relative abundance data, we estimated the absolute abundance of each MAG by converting sequence coverage depths into absolute genome copy numbers using MGCalibrator [58]. This spike-in-free methodology employs a DNA mass-calibration framework previously applied to environmental metagenomes [58–60]. The framework relies on sample-specific calibration scaling factors, computed as the ratio of total extracted DNA mass to the DNA mass of the fraction actually sequenced. The scaling factors are then multiplied by the contig-specific coverage depth to yield calibrated absolute abundances. This calibration framework has been robustly validated to show exceptional correlation (*r*^2^ = 0.96 to 0.98) with absolute qPCR quantification [58]. To generate the underlying coverage profiles, genome coverages in each sample were calculated with CoverM in genome mode using Strobealign for read mapping [parameters: genome -p strobealign -m mean --bam-file-cache-directory]. MGCalibrator was then run using these sample-specific BAM files, filtered to a minimum read mapping identity of 95% [parameters: --filter_percent 95].

### Functional analysis of metagenome-assembled genomes

Protein sequences were predicted for each MAG individually using Prodigal v2.6.3[61] in single genome mode, which was parallelised across 48 CPU cores using GNU parallel [62]. This resulted in a total dataset of 2,370,402 MAG protein sequences. We used the protein structure-guided functional profiling tool EcoFoldDB v2.1.0 [63] [parameters: --gpu 1 –qcov 0.7 --tcov 0.7] to identify genes linked to key ecological traits. EcoFoldDB profiling targeted genes involved in plant-microbe interactions, carbon fixation, trace gas oxidation, as well as nitrogen, phosphorus, and sulphur cycling. To convert these gene-level annotations to genome-level trait profiles, we developed a custom Python script *ecofolddb-genome-trait-matrix.py* (https://github.com/timghaly/EcoFoldDB-genome-trait-matrix). For each genome, the script scores a given ecological trait or pathway as present only if a minimal set of essential core genes for that function are detected. The defining minimal gene configurations used for each trait are listed in Supplementary Table S4. We also screened MAGs for gene clusters encoding iron acquisition, storage and redox cycling functions using FeGenie v1.2 [64]. MAGs belonging to the genus *Trinickia*, were further annotated with DRAM v1.5.0 [65], using the *annotate* and *distill* subcommands with default settings, and run_dbCAN v5.1.2 [66–68] *easy_substrate* command for predicting target substrates of CAZyme gene clusters.

### Statistical analyses

All statistical analyses were conducted in R [69]. To control the false discovery rate (FDR) arising from multiple hypothesis testing, all *p*-values were adjusted using the Benjamini-Hochberg method [70], where appropriate.

To identify metabolic pathways with divergent trajectories between treatments, the MaAsLin2 v1.4.0 R package [71] was used to analyse community-level HUMAnN3 pathway relative abundances. Separate linear models were fit for the drought period (T0-T3) and the recovery period (T3-T6, with T3 re-zeroed as the recovery baseline). Each model used *Treatment*, *Time*, and their interaction (*Treatment* × *Time*) as fixed effects. A third model tested *Treatment* alone at the final time point (T6) to assess whether any drought-associated legacy persisted at the recovery endpoint. All MaAsLin2 models were fit using a log-transformation with no additional normalisation, excluding pathways present in fewer than 10% of samples. Pathways exhibiting a significant *Treatment* × *Time* interaction effect during the drought period (MaAsLin2-recommened threshold: *q*-value < 0.25) were subsequently extracted from the recovery period (T3-T6) and endpoint (T6) models to track their trajectories following the drought period.

To identify putative taxa driving observed pathway enrichments, we tested for correlations between drought-responsive pathways and individual OTUs. First, to select a single ribosomal marker gene as a representative OTU classification, we compared Shannon diversity indices calculated from each individual ribosomal marker against the cross-marker Shannon average (Supplementary Fig. S2). The S5 ribosomal protein marker had the lowest root-mean-square error (RMSE) relative to the average, and was thus selected as the representative OTU classification for all subsequent OTU-based correlation analyses. Associations between S5 OTU abundances and enriched pathways were identified using Spearman rank correlations.

Absolute MAG abundances (MGCalibrator-adjusted depths) were used to calculate a per-genome log_2_ fold-change (FC) at each time point between treatment (drought/rewetting) and control samples. MAGs were considered to have increased under treatment conditions if their log_2_FC > 2 at any time point, or decreased if their log_2_FC < −2. To compare genome-level drought responses between rhizosphere and bulk soil, the proportions of drought-increased and decreased genomes were compared between rhizosphere and bulk soil at each time point using two-proportion *z*-tests.

To identify functional traits associated with these drought responses in the rhizosphere, we coupled non-metric multidimensional scaling (NMDS) ordination of absolute genome abundances with functional trait vector fitting. The underlying NMDS ordination was performed using the *metaMDS* function form the vegan R package based on Bray-Curtis dissimilarities of MAG absolute abundance profiles. MAGs from the obligate symbiotic phyla Patescibacteriota (epibionts) and Babelota (intracellular parasites) were excluded from this analysis since their abundances are intrinsically host-dependent. Functional traits (with non-zero variance) were then fitted as vectors onto the NMDS ordination space using the *envfit* function from the vegan R package with 999 permutations. For traits significantly correlated with the ordination space (FDR-adjusted *p* < 0.05), Fisher’s exact tests were used to determine whether the proportion of genomes carrying each trait differed significantly between the drought-increased and decreased response cohorts.

To identify putative bacterial or archaeal hosts of Patescibacteriota MAGs (also known as the Candidate Phyla Radiation or CPR), we tested for co-abundance between Patescibacteriota MAG depths and S5 OTU abundances using the Hmisc v4.6.0 R package [72]. Spearman correlations were computed for all Patescibacteriota MAG-OTU pairs. MAG-OTU pairs with a positive correlation (Spearman’s ρ ≥ 0.7) and an FDR-adjusted *p*-value < 0.05 were retained as high-confidence putative host associations. Similarly, to identify putative eukaryotic hosts for the intracellular parasitic phylum Babelota, we evaluated co-abundance relationships between Babelota MAGs and microeukaryotic species profiled with EukDetect. Pairwise Spearman rank correlations were calculated for all MAG–eukaryote pairs.

To evaluate the association between the genus, *Trinickia*, and plant host biomass during drought, *Trinickia* relative abundances (derived from SingleM genus profiles) were compared against above- and below-ground plant dry weights for drought-phase rhizosphere samples (T1 –T3). To account for the confounding effects of general plant growth over time, we analysed time-adjusted residuals for each variable. First, natural log-transformed *Trinickia* relative abundance values and plant dry weights were separately regressed against sampling time point in linear models. The resulting residuals, representing variation in *Trinickia* abundance and plant biomass traits independent of time, were then correlated using Spearman rank correlations to test for any underlying associations.

## Results and Discussion

### Decoupled taxonomic and functional diversity trajectories across compartments during drought and rewetting

Here, we examined the impact of drought and subsequent rewetting on soybean microbiomes using a controlled glasshouse experiment, coupled with deep metagenomic sequencing. We found that sustained drought drove an increasing divergence in rhizosphere community composition and functional potential relative to controls (Fig. 1). The drought-driven divergence was characterised by a decrease in taxonomic diversity, alongside a concurrent increase in functional diversity and average prokaryote genome size (Fig. 1). These patterns suggest that while environmental filtering under sustained water deficit lowers rhizospheric microbial diversity, it selects specialised taxa with larger genomes, potentially necessary to encode the expanded functional repertoire required to survive drought stress. Upon rewetting, both the taxonomic and functional composition of the rhizosphere trended back toward the baseline (control) state. However, while functional diversity returned close to control levels, average prokaryotic genome size persisted at a slightly elevated level through to the final sampling time point (Fig. 1). The precise drivers of this lingering genome size footprint remain unclear, particularly given the apparent secondary downward trend in taxonomic diversity at the final time point (T6). These minor late-stage adjustments might represent transient successional fine-tuning as the community re-establishes steady-state interactions with the host.

**Fig. 1.**
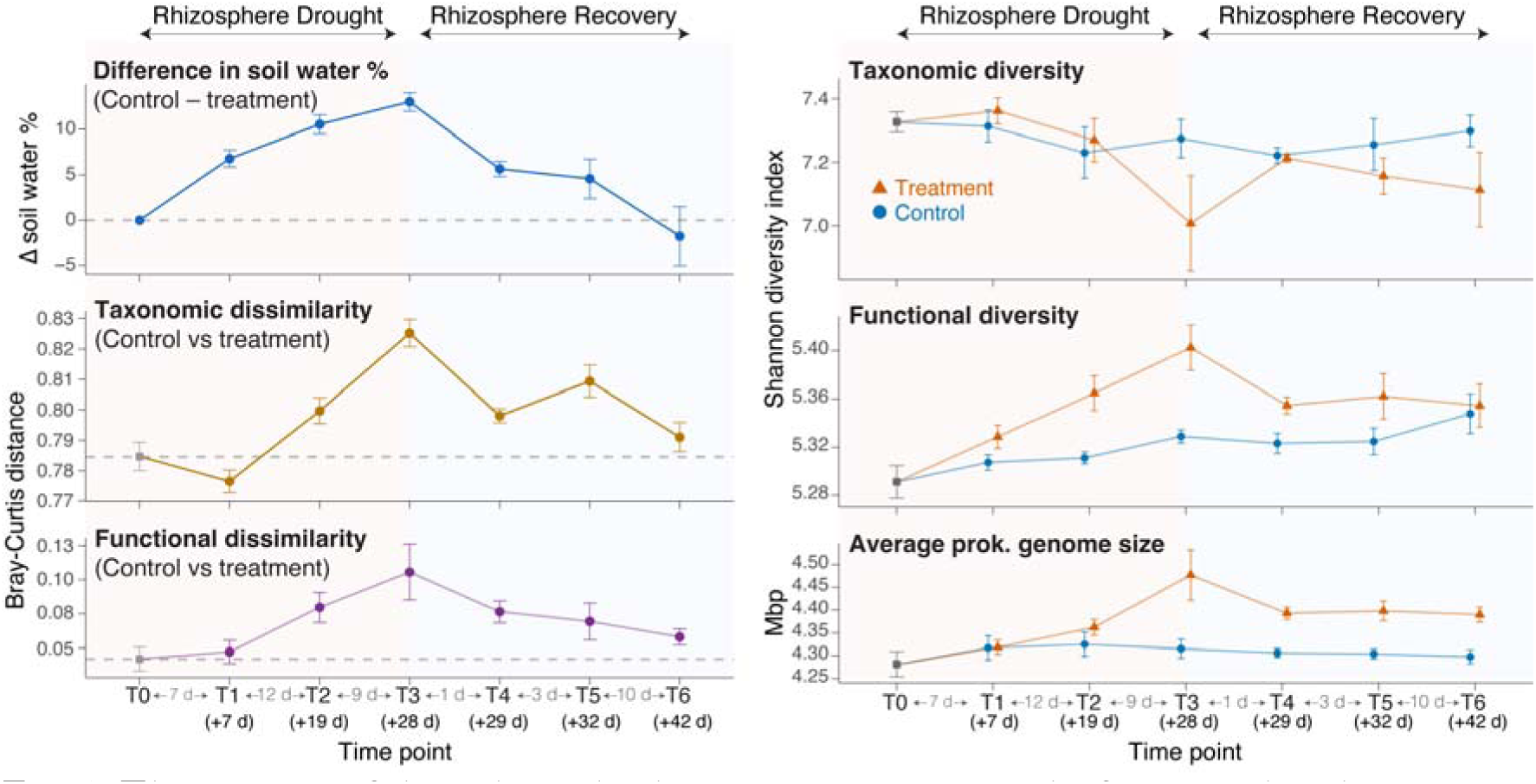
The impacts of drought and subsequent recovery on the functional and taxonomic diversity within the rhizosphere. The shared x-axes indicate sampling time points (T0–T6), with the number of days since commencement of drought treatment shown in parentheses, and the number of days between consecutive sampling points displayed between the tick marks. Left panels illustrate the progression of physical and community-level divergence over time, displaying the mean difference in gravimetric soil water content (GWC, %) between control and drought treatments (top), as well as taxonomic (middle) and functional (bottom) beta-diversity distances (Bray-Curtis dissimilarity) between control and treatment rhizosphere samples. Right panels show alpha-diversity trajectories for treatment (orange triangles) and control (blue circles) lines, showing taxonomic Shannon diversity (top), functional Shannon diversity (middle), and estimated average prokaryotic genome size (bottom) for rhizosphere samples. All error bars represent ± 1 standard error (S.E.) of the mean. Taxonomic alpha- and beta-diversity metrics are calculated from the average of 14 universal ribosomal protein markers (see methods), while functional diversity metrics were calculated from HUMAnN3-profiled pathways.

We also found that proximity to plant roots fundamentally reshaped post-rewetting dynamics, resulting in divergent recovery trajectories between the rhizosphere and bulk soil microenvironments. Functional diversity followed broadly consistent trends across drought and recovery in both compartments, but taxonomic recovery paths diverged sharply: rhizosphere composition trended back toward baseline, whereas bulk soil remained highly dissimilar and variable (Supplementary Fig. S3), pointing to a more persistent drought legacy effect in the absence of root-mediated recolonisation support.

Consistent with the expanded genomic capacity in the rhizosphere under drought (Fig. 1), a community-wide, reads-based analysis revealed a clear enrichment of multiple metabolic pathways (Supplementary Fig. S4). This functional profile was dominated by strategies for surviving osmotic stress, together with mechanisms for scavenging plant-derived carbon and organic nitrogen. For example, osmoprotection pathways (e.g., ectoine biosynthesis) were enriched alongside cell envelope maintenance functions (e.g., lipid, fatty acid, and peptidoglycan biosynthesis and recycling) that preserve structural integrity under drought-induced matric tension [73]. Resource-acquisition potential showed a parallel shift toward organic nutrient pools, marked by enrichment of amino-acid-catabolism pathways for nitrogen scavenging and a broader expansion of carbon-harvesting capacity. This was most pronounced for pathways degrading plant-stress-induced exudate components (e.g., *myo*-inositol, D-galacturonate, and D-glucuronate [74–76]), and the breakdown of complex aromatic compounds (e.g., biphenyl and picolinate degradation), which plants also accumulate under drought stress [77–79]. Upon rewetting, these enriched pathways steadily declined back toward baseline levels (Supplementary Fig. S4). Correlation analysis indicated that these community-wide functional shifts were likely underpinned by a narrow set of taxa. These largely belonged to the family Burkholderiaceae, particularly, the genus *Trinickia*, which showed widespread and strong positive correlations across almost the entire suite of drought-enriched pathways (Supplementary Fig. S4). This suggests that the community-wide functional shifts under drought are strongly underpinned by a highly specific, narrow group of responsive taxa.

Conversely, only three pathways were found to be depleted under drought, including chlorophyllide *a*, thiamine, and fungal flavin biosynthesis. Specifically, the decline in chlorophyllide *a*, and fungal flavin pathways points to a loss of phototrophs (i.e., cyanobacteria/microalgae) and fungi. Direct profiling of the fungal communities themselves yielded no clear successional trends related to drought, likely because the detection limits of shotgun metagenomics resulted in a low number of uniquely resolved fungal species, with an average of just 12 species per sample (Supplementary Fig. S5). In parallel, the depletion of thiamine biosynthesis suggests that the drought-selected taxa rely more heavily on thiamine salvage over biosynthesis. Like the enriched functions, the depleted pathways returned to control levels during the recovery phase (Supplementary Fig. S4).

### Absolute genome abundance dynamics reveal rhizosphere-specific responses to drought

Our genome binning workflow yielded 728 species-level metagenome-assembled genomes (MAGs) meeting the high-purity criteria of <5% contamination and >50% completeness (median completeness = 71.56%, median contamination = 1.08%; Supplementary Table S3). The resulting MAG set captured a substantial proportion of the sequencing data, recruiting an average of 62.15% (IQR: 61.83% –62.93%) of total metagenomic reads per sample with a 95% identity threshold (the widely accepted consensus for prokaryotic species delineation [80]). When focusing exclusively on the prokaryotic fraction of each sample, read mapping accounted for an average of 82.84% (IQR: 82.31% –83.85%) of the prokaryotic community, demonstrating that most of the dominant species have genomic representation in our MAG set. This level of MAG recovery substantially exceeds what is typically reported for genome-resolved metagenomics of rhizosphere communities. For example, in one of the most comprehensive genome-resolved studies examining root-microbiome drought responses to date, only 55 MAGs, representing just 4.8% of sequencing reads, were recovered [14].

Additionally, by using a DNA-mass-calibration approach [58], we converted MAG relative abundances to inferred absolute abundances. This allowed us to overcome the inherent limitations of relative abundance data, where proportional shifts mask whether a population is truly increasing, decreasing, or staying the same. Thus, by tracking the absolute abundances of each genome independently, alongside an exceptionally high level of genomic recovery, we could perform a robust, genome-resolved assessment of community dynamics throughout drought and rewetting phases.

We found that a phylogenetically diverse suite of taxa sharply declined during drought in the rhizosphere, with the most severe decreases observed across genomes belonging to Pseudomonadota (formerly Proteobacteria), Chlamydiota, Bacteroidota, and Gemmatimonadota (Fig. 2a). Conversely, a much narrower diversity of taxa increased in the rhizosphere under drought, heavily dominated by *Trinickia* spp. (Fig. 2a), which mirrors findings from our reads-based analysis (Supplementary Fig. S4). Interestingly, the proliferation of these drought-responsive genomes occurred in two distinct waves, revealing clear successional dynamics as water deficit progressed. Strikingly, rewetting triggered an extremely rapid recovery, with the rhizosphere genome abundances returning to baseline within 24 hours (Fig. 2a), underscoring the resilience of the root-associated microbiota. We also identified host-dependent phyla such as Babelota (intracellular protist parasites [81]) and Patescibacteriota (obligate bacterial epibionts [82]) as highly responsive, comprising individual genomes that either sharply increased or decreased during drought. Because their fitness is intrinsically tied to host survival, these shifts likely reflect indirect drought impacts on their respective hosts. While co-abundance correlations failed to identify candidate hosts for Babelota, several putative bacterial hosts for Patescibacteriota were resolved (Supplementary Fig. S6). Specifically, drought-enriched Patescibacteriota strongly correlated with *Limisphaera*, Burkholderiaceae, and *Mucilaginibacter*, whereas drought-depleted genomes tracked with a single methionine-auxotrophic species, *Lysobacter auxotrophicus*.

**Fig. 2.**
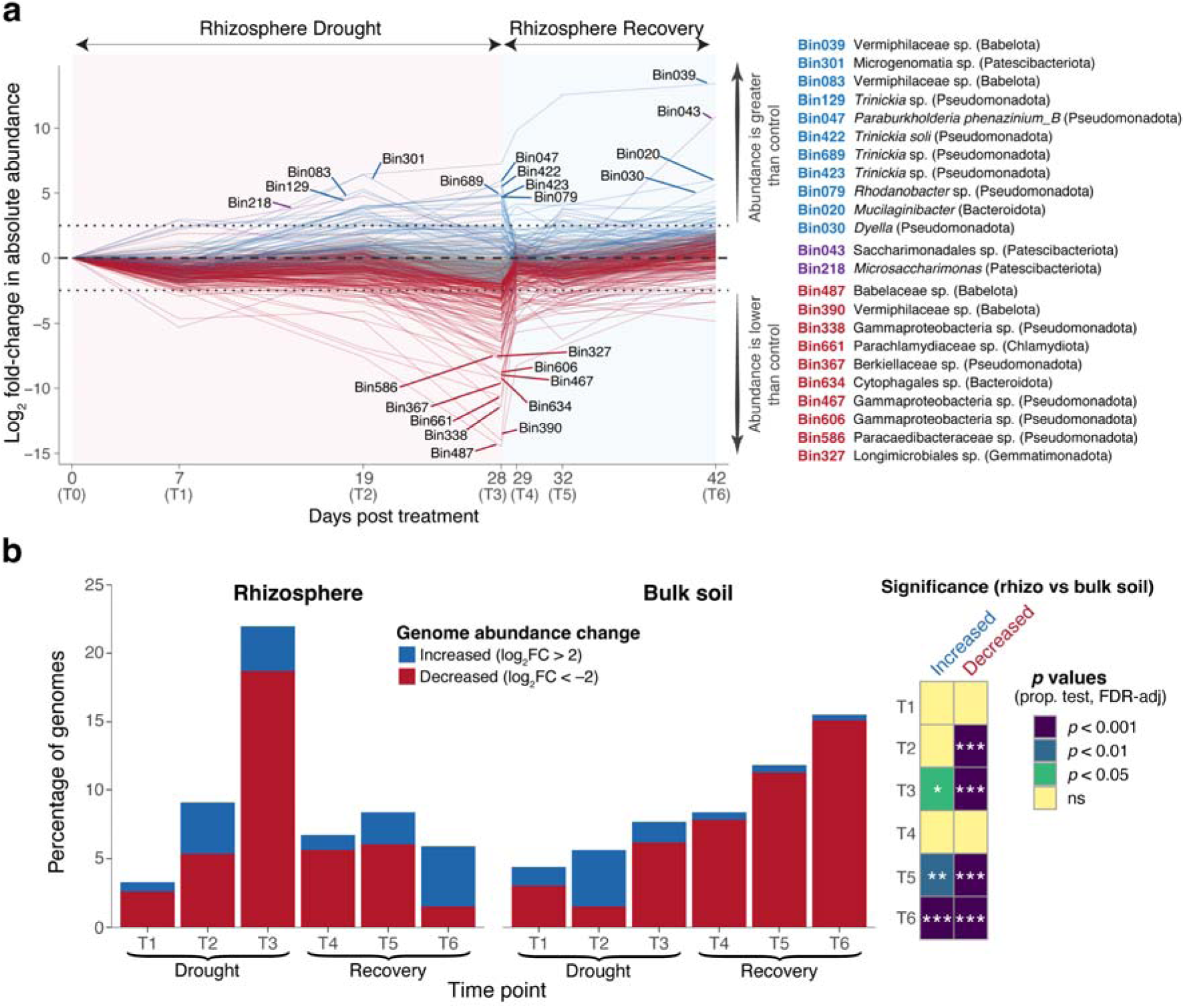
Metagenome-assembled genome (MAG) abundance shifts and rhizosphere-specific responses to drought. (**a**) Temporal trajectories of MAG absolute abundances across the experimental timeline. Blue lines denote genomes that increased in abundance compared to control samples at any given time point (log_2_FC > 2), red lines denote genomes that decreased in abundance (log_2_FC < −2), and purple lines denote genomes that crossed both thresholds at different points during the experiment. The highest-responding MAGs are labelled with their respective lowest taxonomic classifications shown on the right margin. Horizontal dotted lines indicate the log_2_FC thresholds of 2 and −2. (**b**) Stacked bar charts comparing genome-level response between the rhizosphere and bulk soil, with bars reflecting the total percentage of genomes that increased (blue) or decreased (red) relative to control samples at each sampling interval. The adjacent heatmap denotes statistical significance in the proportion of responsive genomes between the rhizosphere and bulk soil compartments at each discrete time point. Asterisks indicate degree of significance (* *p* < 0.05, ** *p* < 0.01, *** *p* < 0.001; ns = non-significant).

We found that genome abundance trajectories differed significantly between rhizosphere and bulk soil compartments (Fig. 2b). Although most genomes did not shift in abundance in either compartment, even at the peak of drought, the rhizosphere harboured a significantly higher proportion of drought-increased and -decreased genomes (Fig. 2b). This suggests that the root-associated environment provides a greater degree of environmental filtering during sustained water deficit, likely driven by stress-induced shifts in plant root exudation and mucilage profiles that are actively enriching specific taxa, while other groups decline [22]. This pattern aligns with observations across distinct plant and cropping systems, where the impact of drought on microbial community composition becomes progressively more pronounced closer to the plant roots [17, 18]. Interestingly, while rhizosphere populations rapidly stabilised toward baseline levels during the recovery period, the bulk soil exhibited a progressive, post-wetting decline in MAG absolute abundances (Fig. 2b), which is also reflected in our reads-based analysis (Supplementary Fig. S4). These findings mirror previous reports of a drought legacy effect in bulk soil that is not seen in the wheat rhizosphere [18]. The bulk soil community thus appears to sustain an accumulative disturbance trajectory that continues after the drought stress is removed.

Here, we hypothesise that rewetting in the bulk soil triggers a successional disturbance rather than a recovery. We propose that this continued disturbance trajectory could be driven by the Birch effect, a well-documented phenomenon in which rewetting of dry soil produces a massive pulse of carbon and nutrients [83]. This sudden water influx forces soil bacteria to rapidly release accumulated osmolytes to survive this osmotic shock, directly inputting labile organic carbon and nitrogen into the soil matrix [84, 85]. Simultaneously, the reconnection of water films enables the diffusion of bioavailable substrates that accumulated during the drought, throughout which, persistent extracellular enzymes continued to generate products that lacked the transport pathways necessary to diffuse [10, 86]. Together, these processes rapidly and substantially alter the chemical profile of the bulk soil post-rewetting, creating a novel successional niche rather than a return to pre-drought conditions. Consequently, fast-growing, copiotrophic taxa can rapidly proliferate to exploit these resources, shifting the community composition further from its baseline state. In contrast, the rhizosphere’s rapid recovery upon rewetting suggests that it is shaped primarily by the influence of root exudates rather than the nutrient pulse of the Birch effect, which is more likely to dominate the comparatively nutrient-limited bulk soil.

Given the healthy root nodules observed on the soybean plants (Supplementary Fig. S7a), we also screened the recovered MAGs for any putative symbiotic rhizobia to examine their dynamics under drought. We recovered a single MAG, *Bradyrhizobium diazoefficiens*, containing essential nitrogen fixation (*nifH*, *nifD*, and *nifK*) and root nodulation (*nodA*, *nodB*, and *nodC*) genes, which shared 99.99% ANI with the Australian commercial soybean inoculant strain CB1809 (Accession: NZ_CP088088.1). Since the seeds were uninoculated, this strain’s higher baseline abundance in the bulk soil relative to the rhizosphere (Supplementary Fig. S7b) suggests that it was already present within the collected topsoil, where commercial rhizobia are known to persist in soils even years after inoculation [87]. We found that *B. diazoefficiens* was highly robust to the effects of drought, regardless of compartment (Supplementary Fig. S7b). This observation extends existing reports of CB1809’s tolerance to desiccation stress during inoculation [88], providing evidence that this resilience also translates to robustness in soil under drought.

### Functional traits associated with genome drought responses in the rhizosphere

To investigate the metabolic and ecological traits driving rhizosphere drought responses, we coupled a non-metric multidimensional scaling (NMDS) ordination of absolute genome abundances with functional trait vector fitting (Fig. 3a). While no conserved traits significantly correlated with drought-decreased genomes, likely due to the broad phylogenetic diversity of this collapsing cohort, a distinct functional signal characterised the drought-increased population in the rhizosphere (Fig. 3a,b). The over-represented traits within this resilient cohort spanned crucial nutrient-scavenging pathways and stress-tolerance mechanisms. Specifically, we observed an enrichment of iron acquisition pathways involved in siderophore synthesis and uptake, alongside haem utilisation as an alternative organic iron source. This was accompanied by heightened capacities for organic phosphorus foraging via phosphatases and carbon-phosphorus (C-P) lyases; nitrogen utilisation through urease and nitrate reduction (NAR); and a comprehensive suite of sulphur-cycling traits spanning organic sulphur mineralisation, sulphite oxidation, and sulphide oxidation. Together, these traits point to an enhanced functional capacity to pivot toward alternative and organic nutrient pools, a critical adaptation when the contraction of soil water films severely restricts the passive diffusion of soluble inorganic forms.

**Fig. 3.**
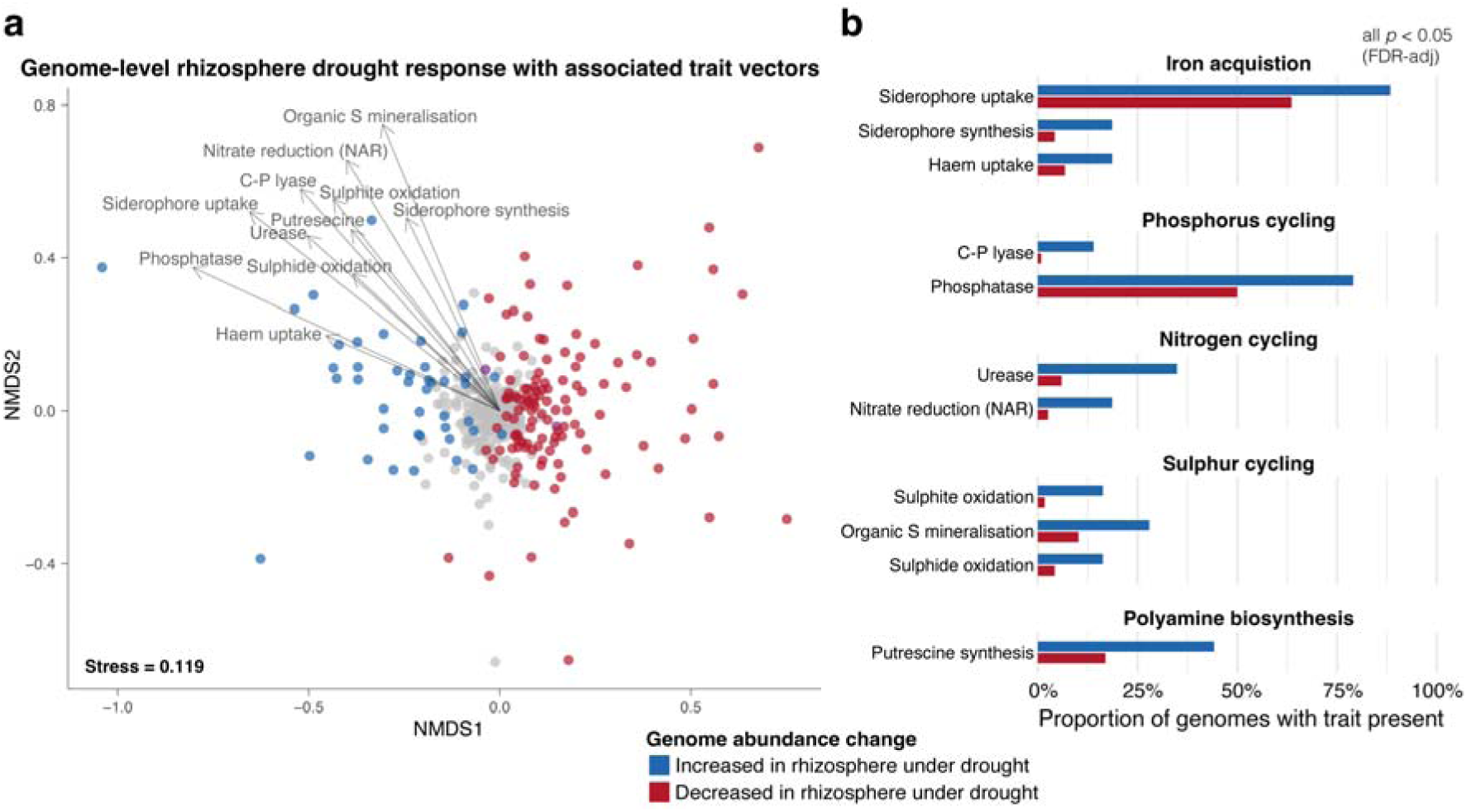
Functional traits associated with genome drought responses in the rhizosphere. (a) Non-metric multidimensional scaling (NMDS) ordination of MAGs based on their absolute abundance profiles in rhizosphere samples. Points represent individual genomes, coloured by their ecological response to drought conditions (Blue: increased in the rhizosphere during drought; Red: decreased; Grey: unchanged relative to controls). Vector overlays represent key ecological traits that significantly correlated with the ordination space. (b) Bar charts displaying the proportion of genomes from each drought-response cohort that encodes each significantly correlated trait. All displayed traits were significantly over-represented in the drought-increased cohort relative to the drought-decreased cohort (Fisher’s exact test, all FDR-adjusted *p* < 0.05).

Beyond nutrient foraging, cellular protection via polyamine biosynthesis was significantly over-represented in the drought-resilient cohort, driven by a sharp elevation in the proportion of genomes encoding putrescine synthesis (Fig. 3b). Polyamines, including putrescine, are well-characterised stress-tolerance molecules that mitigate osmotic and oxidative damage in both bacteria and plants by stabilising cellular membranes and scavenging reactive oxygen species [7, 8]. Thus, in the rhizosphere, microbially derived putrescine may serve a dual ecological function, buffering local osmotic stress for the bacterial producer, and act as a cross-kingdom signalling molecule that directly modulates host plant physiological defences under water deficit.

### Genomic traits of Trinickia proliferation and plant growth promotion under drought

Given that *Trinickia* species appeared to be the dominant drivers of both structural and functional shifts within the rhizosphere during drought, we performed an in-depth functional profiling of the *Trinickia* MAGs in our dataset (*n* = 11 species-level MAGs). From this, we generated a consensus metabolic characterisation of the enriched *Trinickia* species, revealing a robust suite of core drought-survival and plant-growth-promoting traits, potentially explaining their increased fitness in the rhizosphere under drought conditions (Fig. 4).

**Fig. 4.**
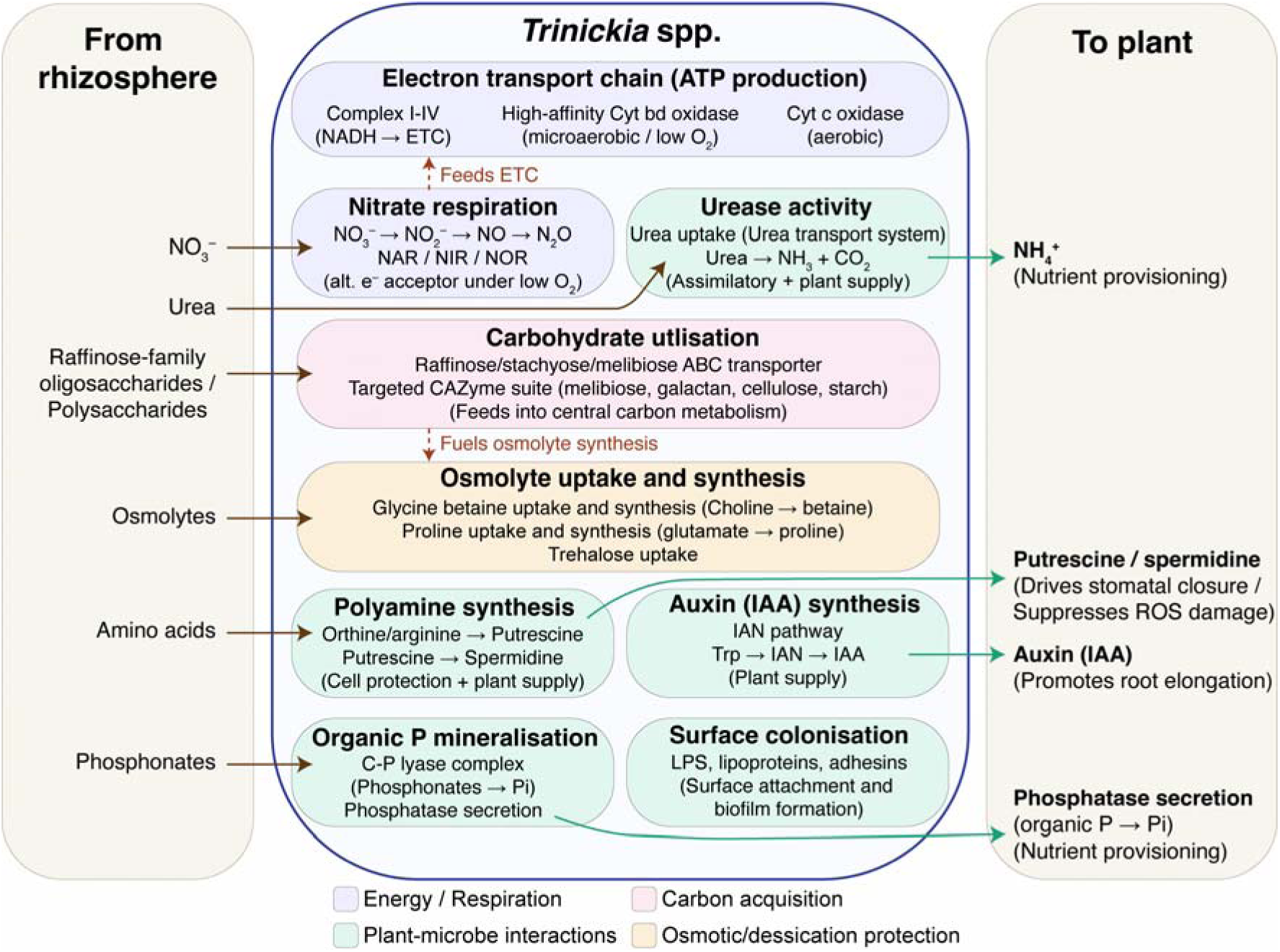
Metabolic model of drought-induced plant-microbe interactions mediated by enriched *Trinickia* spp. Brown arrows trace substrate uptake from the rhizosphere (left), through internal metabolic pathways (middle), and their subsequent output (green arrows) as plant-available nutrients, plant signalling molecules, or phytohormones (right). Metabolic modules are coloured by functional category: energy/respiration (light purple), carbon acquisition (pink), plant-microbe interactions (green), and osmotic/desiccation protection (light orange). Specialised adaptations for growth under drought conditions in the rhizosphere are highlighted. These include uptake of drought-induced root exudates (e.g., raffinose-family oligosaccharides), and microaerobic survival where alternative electron acceptance (nitrate respiration) complements high-affinity oxygen scavenging (cytochrome *bd* oxidase) for energy production when oxygen becomes restricted in drought-compacted soils. *Trinickia* additionally encode pathways involved in plant-growth-promoting activities, including nutrient provisioning, polyamine synthesis (putrescine/spermidine) and auxin production (indole-3-acetic acid; IAA). Functional annotations for the Trinickia genomes are supplied as Supplementary Table S5.

Core drought-survival traits within the recovered *Trinickia* genomes centred around osmolyte uptake and synthesis, alongside energy production under fluctuating oxygen gradients, a hallmark of drying soils in which water retreats into finer pores and creates localised pockets of both high and low O_2_ diffusion [89]. To maintain ATP production under variable oxygen levels, *Trinickia* encode a flexible dual-terminal electron transport chain featuring a high-affinity cytochrome *bd* oxidase alongside aerobic cytochrome c oxidases (Fig. 4). This high-affinity system facilitates microaerobic respiration within oxygen-limited micro-niches. This is complemented by a complete nitrate respiration pathway (Fig. 4) that uses nitrate (NO_3_^−^) as an alternative electron acceptor under oxygen limitation [90]. Additionally, to mitigate osmotic and desiccation stress under water deficit, *Trinickia* possess pathways for both the exogenous uptake and *de novo* biosynthesis of the osmolytes, glycine betaine (via choline precursors) and proline (via glutamate pathways), alongside specialised ABC transporters for trehalose uptake and cycling, providing robust protection against desiccation (Fig. 4).

*Trinickia* also encode key plant growth-promoting traits critical for buffering plant hosts against drought stress. For instance, drought physically hinders the mass flow and diffusion of inorganic nutrients, making nitrogen and phosphorus highly immobile and driving plant nutrient deficiencies [6]. Here, *Trinickia* can potentially assist in plant nutrient provisioning by mineralising organic forms of phosphorus. Specifically, *Trinickia* encode C-P lyase as well as secreted phosphatases that mineralise organic phosphonates into bioavailable orthophosphate (P*_i_*) directly at the root interface (Fig. 4). Similarly, organic nitrogen is mineralised via *Trinickia*-encoded urease activity, converting urea into available ammonium (NH_4_^+^). Consequently, *Trinickia* possess the functional potential to enhance P*_i_* and NH ^+^ availability for both itself, and its plant host precisely when the physical mobilisation of inorganic forms is most restricted. Complementing this nutrient provisioning, *Trinickia* encode pathways for the production of the auxin phytohormone, indole-3-acetic acid (IAA), and the polyamines, putrescine and spermidine (Fig. 4). *Trinickia*’s biosynthesis of IAA can drive root elongation and architectural modifications that help plants survive during prolonged dry spells [2–4]. Specifically, these IAA-induced shifts can help optimise water foraging deeper in the soil profile as well as stimulating lateral root branching and root hair density to expand the overall absorptive surface area of the root system [4, 5]. Meanwhile, polyamines can function as cross-kingdom signalling molecules that further mitigate host stress [5]. Putrescine triggers leaf stomatal closure, which helps minimise transpirational water loss, while spermidine can suppress host tissue damage from reactive oxygen species (ROS) that spike under water deficit [91–94].

The striking enrichment of *Trinickia* within the rhizosphere, which is not observed in the surrounding bulk soil (Supplementary Fig. S8), appears to be mediated by a targeted capacity to exploit plant stress exudation profiles (Fig. 4). Under water deficit, plants frequently over-produce raffinose-family oligosaccharides (RFOs) as systemic osmoprotectants [95–97], while also altering the composition of their exuded, gelatinous root mucilage [19, 21]. *Trinickia* appear to leverage this stress-induced resource shift by encoding a specialised RFO transporter system (Fig. 4). This uptake machinery is paired with a targeted suite of carbohydrate-active enzymes (CAZymes) tailored to metabolise soluble RFO breakdown intermediates like melibiose [98], as well as galactan, a major polysaccharide component of root mucilage [99]. Channelling these host-derived carbon streams into central carbon metabolism likely allows *Trinickia* to sustain rapid growth during drought and meet the energetic demands of other metabolic functions, such as osmolyte synthesis. Collectively, these traits likely give *Trinickia* a clear competitive advantage in the rhizosphere during water deficit, while also promoting host plant survival through a mutualistic feedback loop.

### Trinickia abundance correlates with host root biomass

Interestingly, we found no significant effects of water deficit on plant host traits, including shoot and root biomass (Supplementary Fig. S9), indicating a highly resilient host response to drought. We therefore tested whether *Trinickia* could be actively buffering the effects of water deficit on the plant host against water deficit by examining which plant traits correlated with its relative abundance. For this, we used a partial regression to separate the influence of *Trinickia* from general plant development across sampling dates (Fig. 5). After accounting for the effect of developmental time, a strong positive correlation was observed between *Trinickia* relative abundance and root biomass (Spearman’s ρ = 0.73, *p* = 0.025). This clearly shows that host plants that harboured a higher proportion of *Trinickia* in their root microbiomes maintained larger root systems relative to their developmental stage. Since, for this specific analysis, we opted to measure *Trinickia* as relative, instead of absolute, abundance, this correlation reflects selective enrichment in larger root systems rather than passive scaling of microbial load with root size. We also found a general trend for increased shoot biomass with *Trinickia* relative abundance, although this was not statistically significant (Supplementary Fig. S10; Spearman’s ρ = 0.43, *p* = 0.25).

**Fig. 5.**
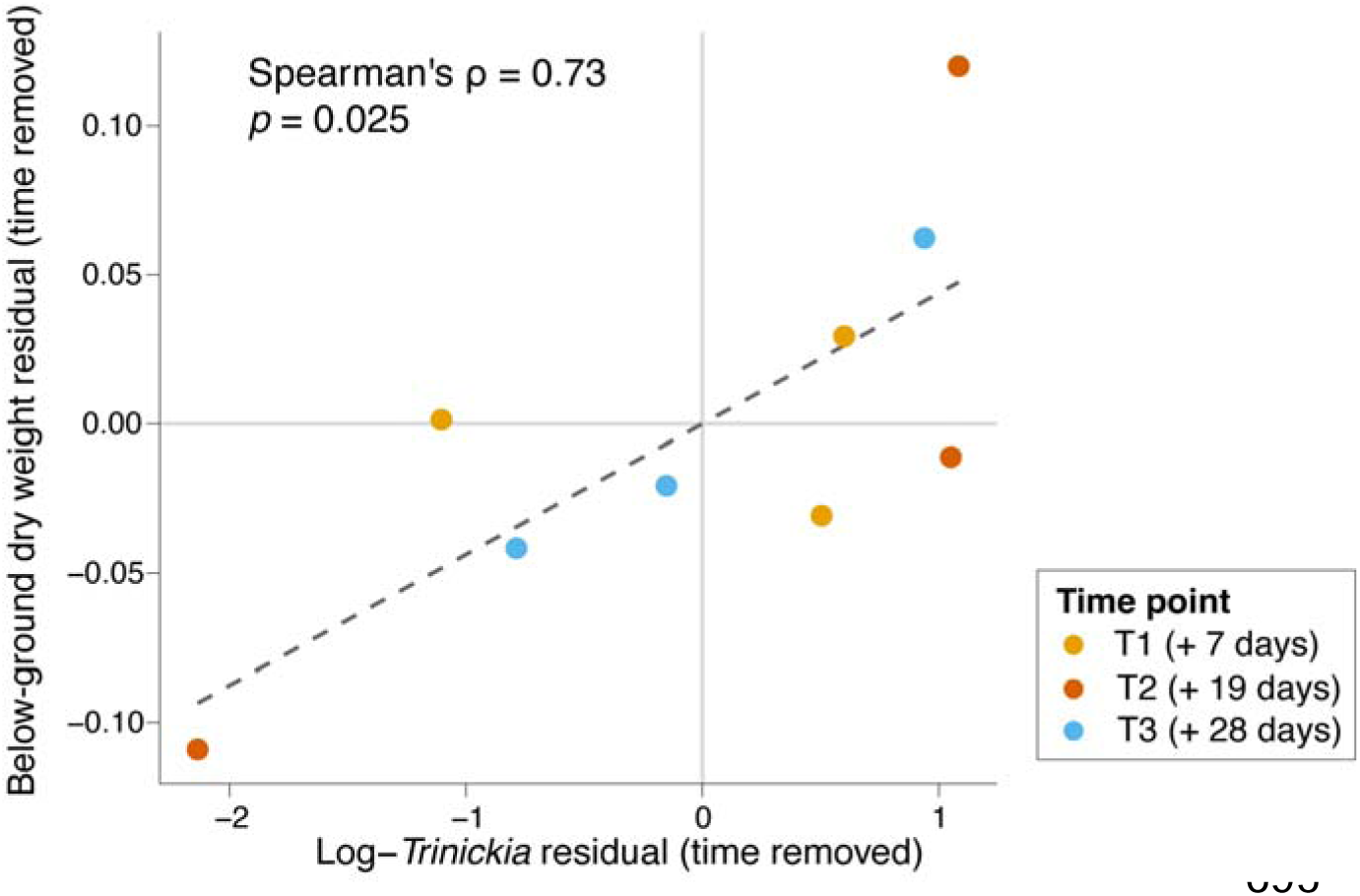
*Trinickia* relative abundance correlates with plant host below-ground weight. Time-adjusted trend between the natural log of *Trinickia* relative abundance and below-ground dry weights, tracked across individual drought sampling time points (T1, T2, and T3). Axes show time-removed residuals indicating a strong, positive underlying relationship between *Trinickia* relative abundance and dry root weight once the confounding effects of general plant growth over time is statistically removed (Spearman’s ρ = 0.73, *p* = 0.025; dashed line indicates linear fit).

These empirical findings support our genomic predictions of plant growth promotion, suggesting that *Trinickia* could indeed be actively contributing to the observed host plant drought resilience. In particular, this root-focused enhancement aligns with *Trinickia*’s capacity for IAA biosynthesis (Fig. 4), known to drive expanded root development. Thus, the drought-induced proliferation of *Trinickia* in the rhizosphere may not merely be a passive response to stress-induced exudation, but might also serve to enhance host root development precisely when soil moisture becomes limiting.

## Conclusion

Here, we show that rhizosphere and bulk soil microbial communities differ fundamentally in how they respond to, and recover from, drought. Resolving these dynamics at genome resolution and in absolute rather than relative abundance terms, we find that drought imposes substantially stronger environmental filtering within the root zone than in bulk soil. This enriched rhizosphere community was dominated by a single, relatively uncharacterised genus, *Trinickia*, whose metabolic profile is geared toward drought survival, nutrient provisioning, and the utilisation of common plant stress-induced exudates. This points to a positive host–*Trinickia* feedback loop under water deficit. Critically, this genomic inference is matched by a direct phenotypic outcome: *Trinickia* abundance correlates strongly with increased host root biomass during active drought, providing empirical support for its role in host drought resilience.

The divergence between compartments becomes even more pronounced after rewetting. Rhizosphere genome abundances recovered to baseline within 24 hours, whereas the bulk soil community continues a progressive trajectory of disturbance. We hypothesise that rewetting acts as a secondary successional disruption in the bulk soil, potentially driven by the Birch effect, which can drastically alter the nutrient profile of the rewetted soil. Together, these findings show that plant roots actively enrich specific beneficial taxa under water deficit and drive the community’s rapid return to its pre-drought state; a recovery not observed in the bulk soil even 14 days after rewetting.

Ultimately, these distinct spatial dynamics frame the rhizosphere as a highly responsive compartment to drought and rewetting conditions. The metabolic adaptation and rapid recovery of root-associated taxa give a clear route to support the host when water and nutrient diffusion are most restricted, while the bulk soil’s prolonged destabilisation exposes the vulnerability of the wider soil matrix. Understanding these distinct spatial trajectories provides a clearer framework for predicting rhizosphere and soil ecosystem resilience to climatic extremes. Beyond this ecological picture, identifying *Trinickia* species as promising candidates for microbiome-based strategies to enhance soybean drought resilience can help inform the rational design of drought-resilient microbiomes.

## Availability of data and material

Metagenomic sequence data generated in this study have been made available via the European Nucleotide Archive (ENA) under study accession PRJEB102647 (BioSamples: SAMEA120513386 –SAMEA120513463).

## Authors’ contributions

T.M.G. conceived and designed the study, secured funding, contributed to experimental work, conducted data analysis, and wrote the original manuscript draft. V.M.J., V.R. and E.C. contributed to laboratory analyses. S.G.T. secured funding and contributed to project design and management. All authors reviewed and edited the final manuscript.

## Acknowledgements

TMG would like to thank Mary, Saoirse and Maebh Ghaly for their loving support.

