## Supplementary Figs. S1-S10 for "Drought and rewetting drive divergent microbial dynamics across soil compartments and reveal *Trinickia* as a key genus for soybean drought resilience"

**This file includes:**

Supplementary Figs. S1-S10

**Other Supplementary Data for this manuscript include the following:**

Supplementary Table S1-S5

**Supplementary Figures**

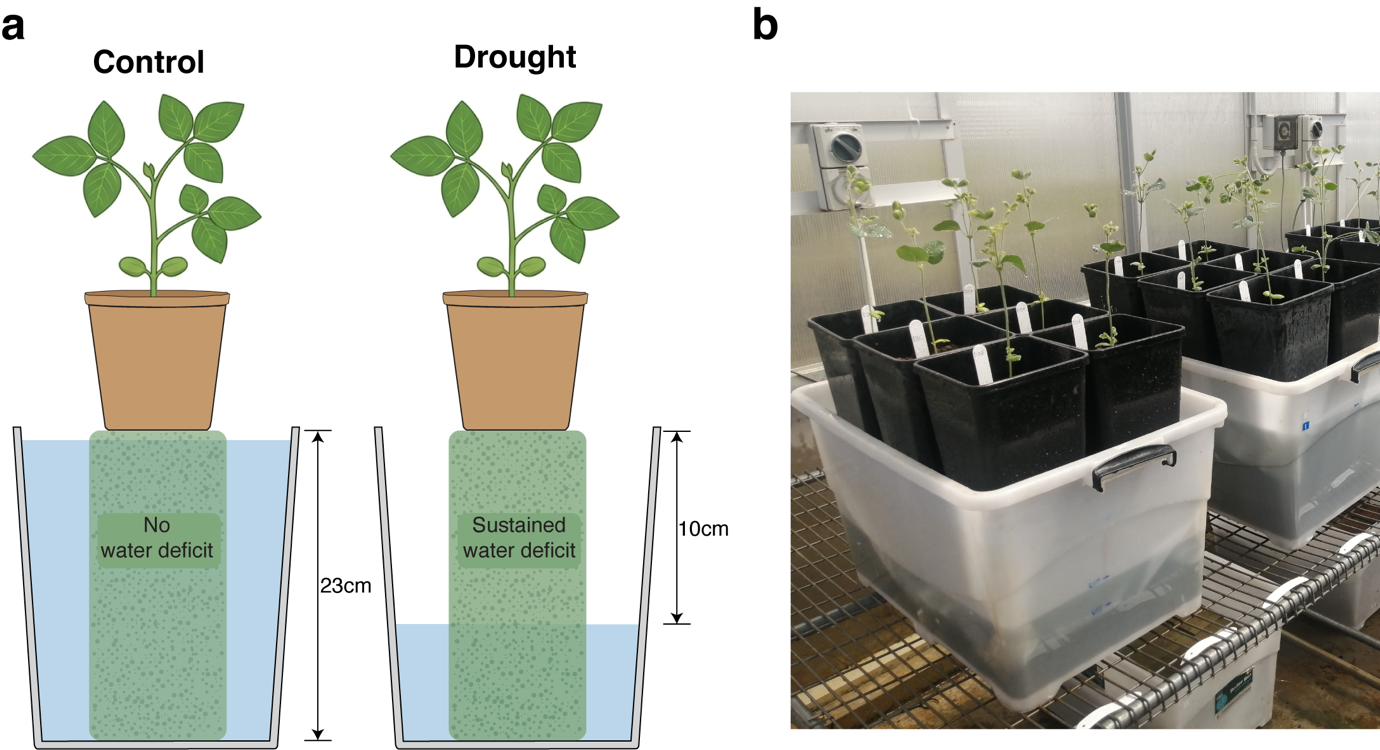
**Fig. S1.** **Experimental setup of plant drought treatment.** (**a**) A diagram of the water deficit method used. Potted plants were placed above a solid column of commercial porous foam (green blocks) inside plastic tubs. Capillary irrigation was used to control soil water content of potted plants, where a greater vertical distance between the base of the pot and the water table produces a more intense drought. The water table level was maintained at the top of the foam blocks for control pots (left), and at a vertical distance of 10 cm for the drought-treated pots (right). (**b**) A photo of the potted soybean plants during the drought phase of the experiment.

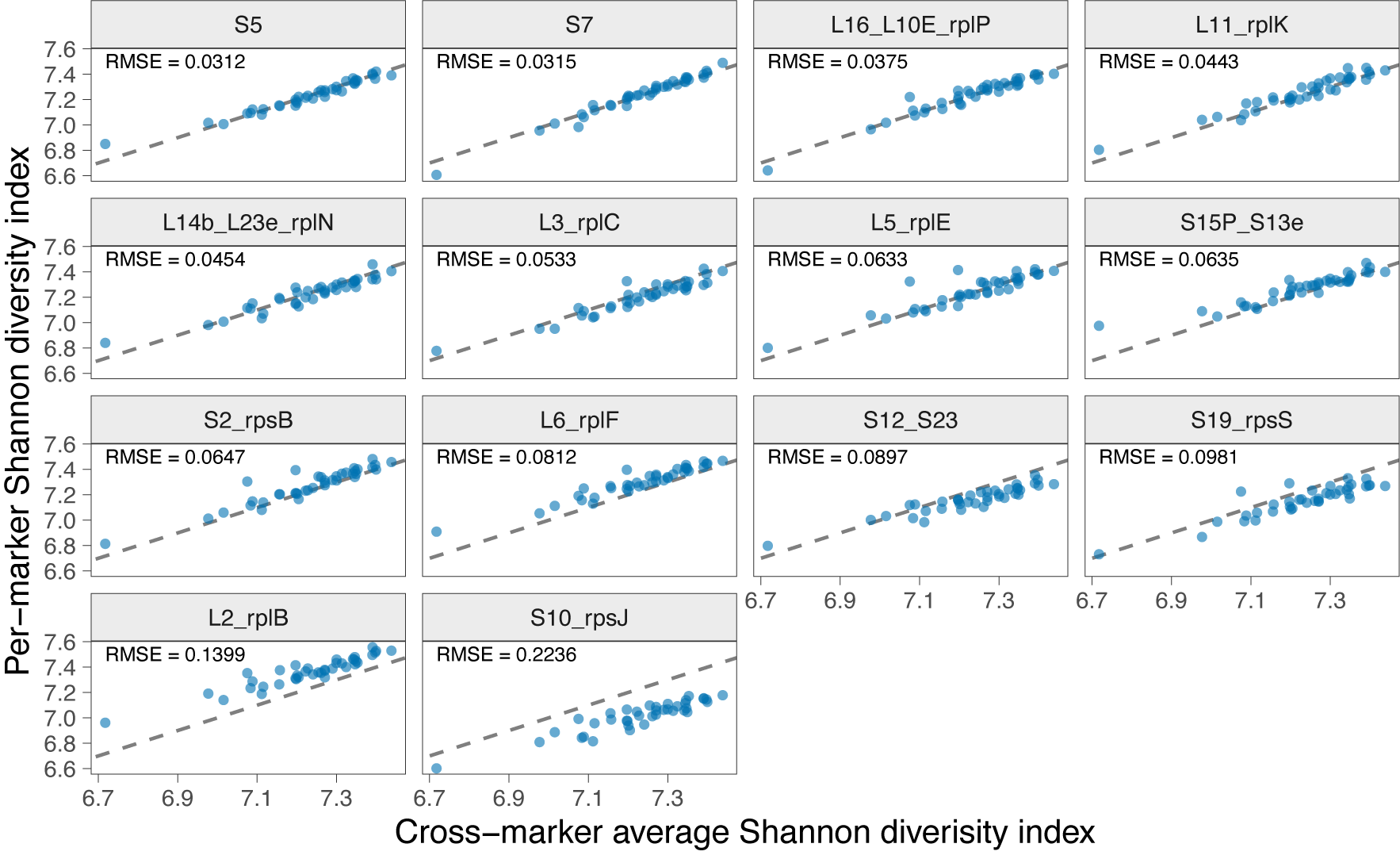

**Fig. S2**. **Selection of single-best representative ribosomal protein marker for OTU correlations.** Per-maker Shannon diversity (y-axis) vs cross-marker Shannon average (x-axis). OTU counts for each of the 14 ribosomal protein markers were generated by SingleM. Each panel represents a different marker, and ordered by lowest root-mean-square error (RMSE) relative to the average (best top-left).

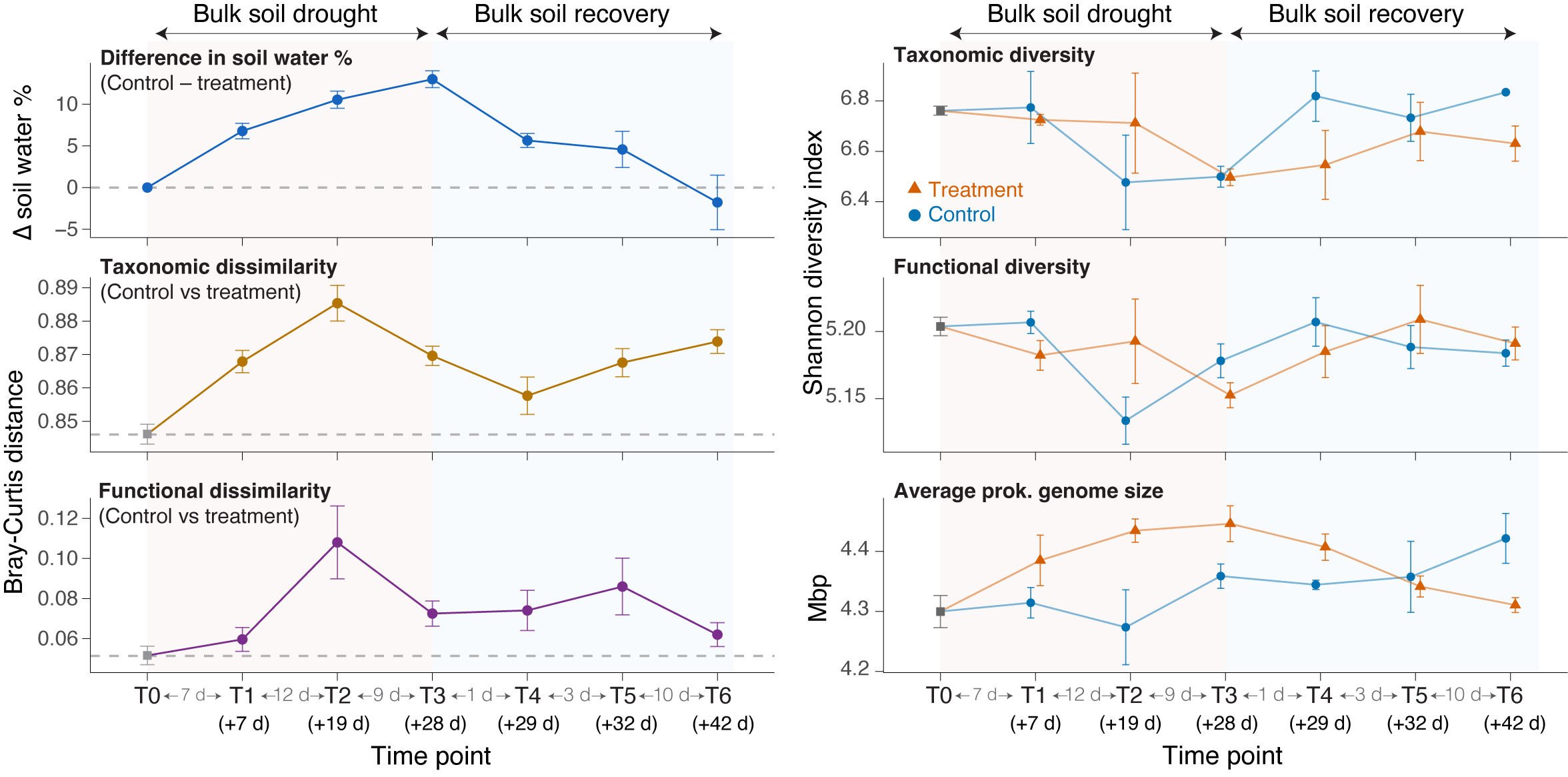

**Fig. S3. The impacts of drought and subsequent recovery on the functional and taxonomic diversity within the bulk soil.** Left panels illustrate the progression of physical and community-level divergence over time, displaying the mean difference (± 1 S.E.) in gravimetric soil water content (GWC, %) between control and drought treatments (top), as well as taxonomic (middle) and functional (bottom) beta-diversity distances (Bray-Curtis dissimilarity) between control and treatment bulk soil samples. Right panels show alpha-diversity trajectories for treatment (orange triangles) and control (blue circles) lines, showing taxonomic Shannon diversity (top), functional Shannon diversity (middle), and estimated average prokaryotic genome size (bottom) for bulk soil samples. Taxonomic alpha- and beta-diversity metrics are calculated from the average of 14 universal ribosomal protein markers (see methods), while functional diversity metrics were calculated from HUMAnN3-profiled pathways.

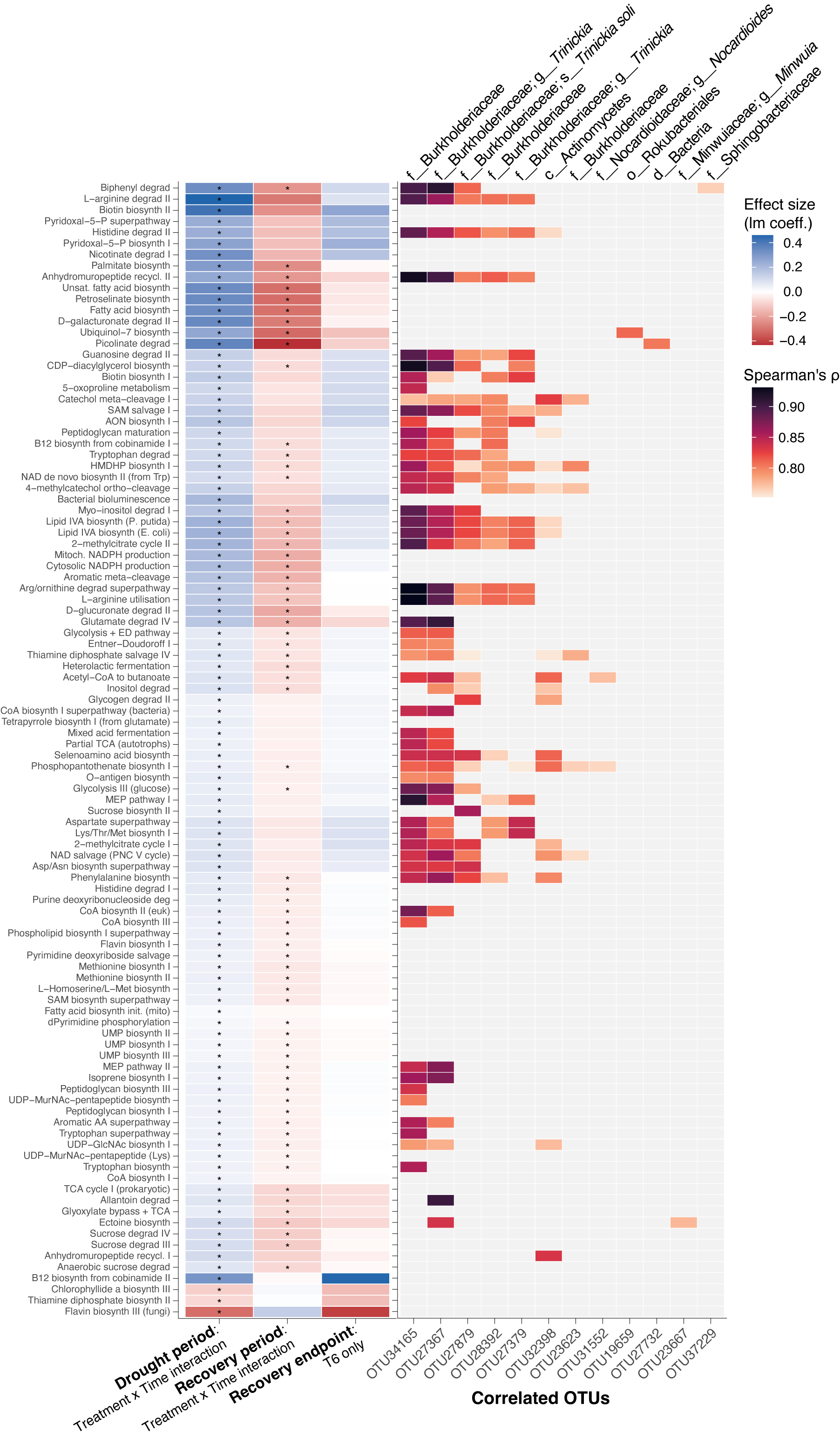

**Fig. S4. Enriched metabolic pathways and correlated OTUs in the rhizosphere under drought.** Left panel heatmap shows HUMAnN3-profiled pathways exhibiting a significant Treatment × Time interaction effect during the drought period (first column: T0 to T3), and their subsequent effects size during the recovery period (second column: T3 to T6) and recovery endpoint (third column: T6 only). Colours indicate effect size (linear model coefficient), while asterisks indicate significance (MaAsLin2 *q*-value < 0.25). Right panel heatmap shows OTUs identified to be significantly correlated with pathways (Spearman’s ρ ≥ 0.7 and FDR-adjusted *p* < 0.05). OTUs are labelled with their lowest SingleM taxonomic classification based on the Genome Taxonomy Database.

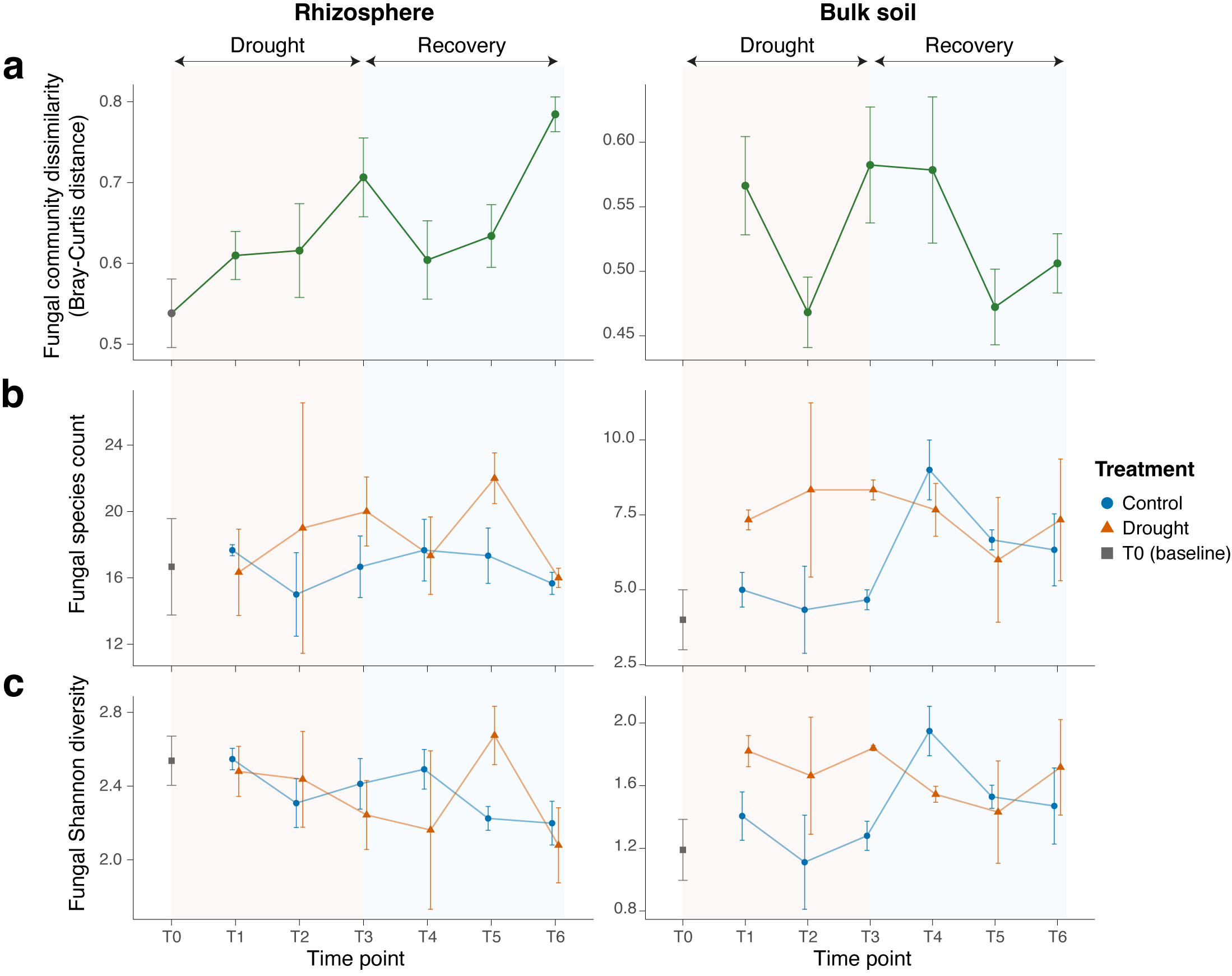

**Fig. S5. The impacts of drought and recovery on fungal diversity within the rhizosphere and bulk soil compartments.** Fungal diversity values are displayed for rhizosphere samples (left panels) and bulk soil samples (right panels), comparing control (blue circles) and treatment (orange triangles) samples for each sampling time point. (**a**) Dissimilarity in fungal community composition (Bray-Curtis distances). (**b**) Observed fungal species counts. (**c**) Shannon diversity values for fungal species.

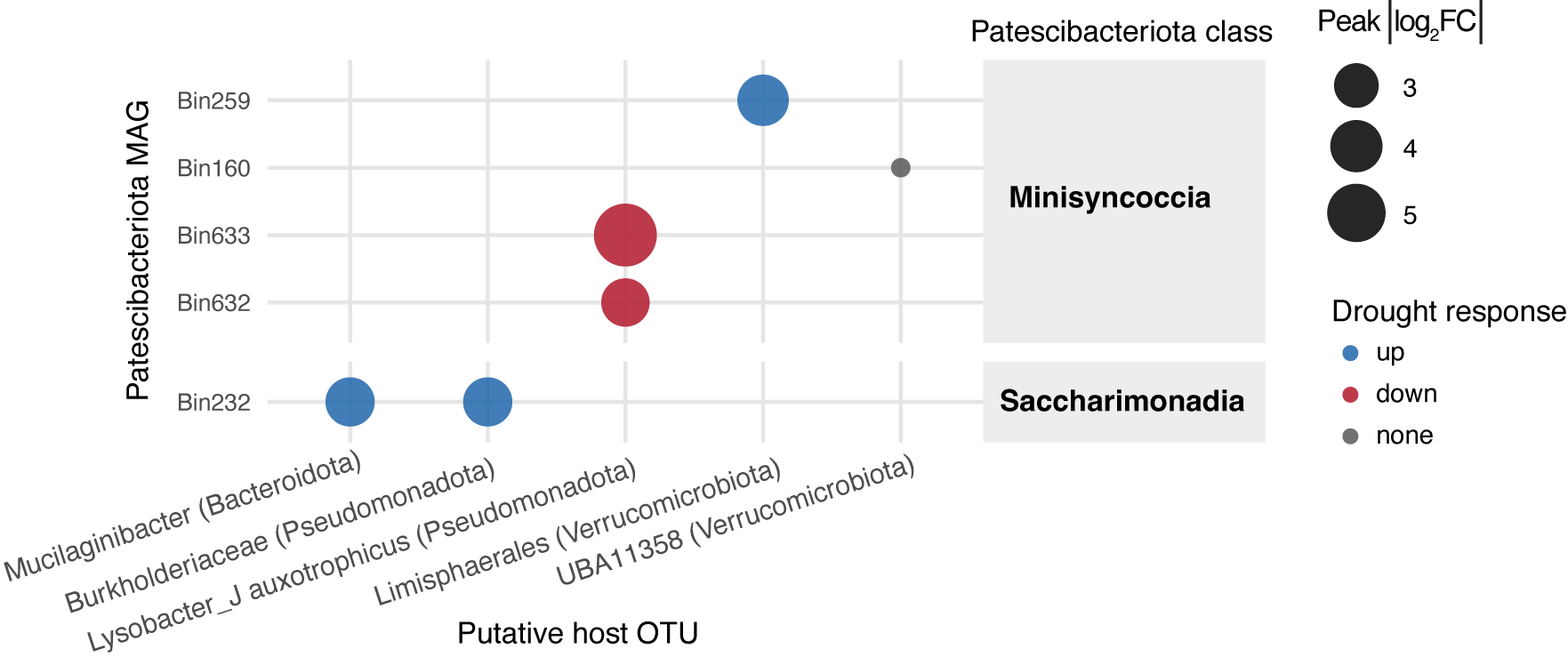

**Fig. S6. Putative host taxa of Patescibacteriota metagenome-assembled genomes.** OTUs identified as putative hosts of Patescibacteriota MAGs via co-abundance analysis (Spearman’s ρ ≥ 0.7 and FDR-adjusted *p* < 0.05). Circle sizes represent the peak absolute log_2_ fold change (FC) in MAG abundance between treatment and control samples at any given time point. Circles are coloured by the MAG’s assigned drought response: drought-increased (blue; log_2_FC > 2), drought-decreased (red; log_2_FC < –2), and neutral MAGs (grey; –2 < log_2_FC < 2).

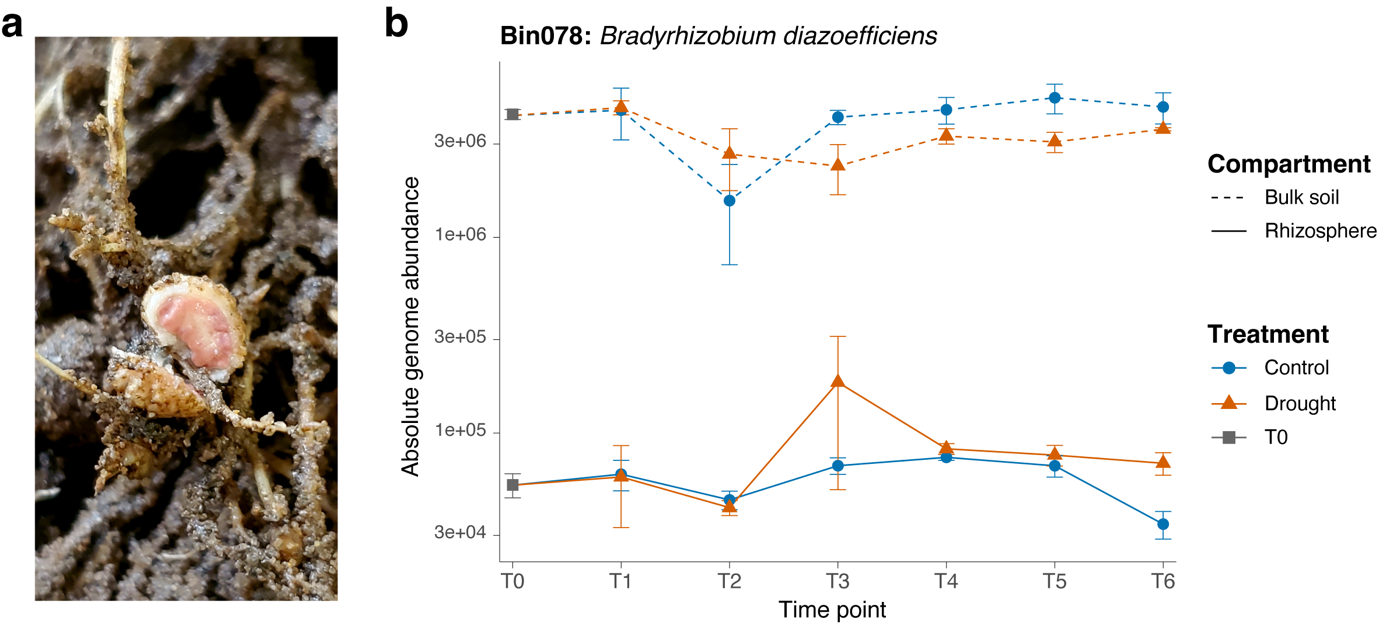

**Fig. S7. Phenotypic evidence of nodulation and absolute genome abundance of identified symbiotic rhizobial metagenome-assembled genome.** (**a**) A dissected soybean root nodule displaying the characteristic pink interior indicative of leghaemoglobin production during active nitrogen fixation. (**b**) A single MAG, *Bradyrhizobium diazoefficiens*, was the only identified MAG to contain nitrogen fixation genes (*nifH*, *nifD* and *nifK*) and root nodulation genes (*nodA*, *nodB* and *nodC*). Points indicate average absolute genome abundance (± 1 S.E.) for *B. diazoefficiens* at each time point for control (blue circles) and drought-treated (orange triangles) samples, and for rhizosphere (solid lines) and bulk soil (dashed lines) compartments. N.B., Rhizosphere soil was collected from intact root systems, and nodules were not independently sampled for DNA sequencing.

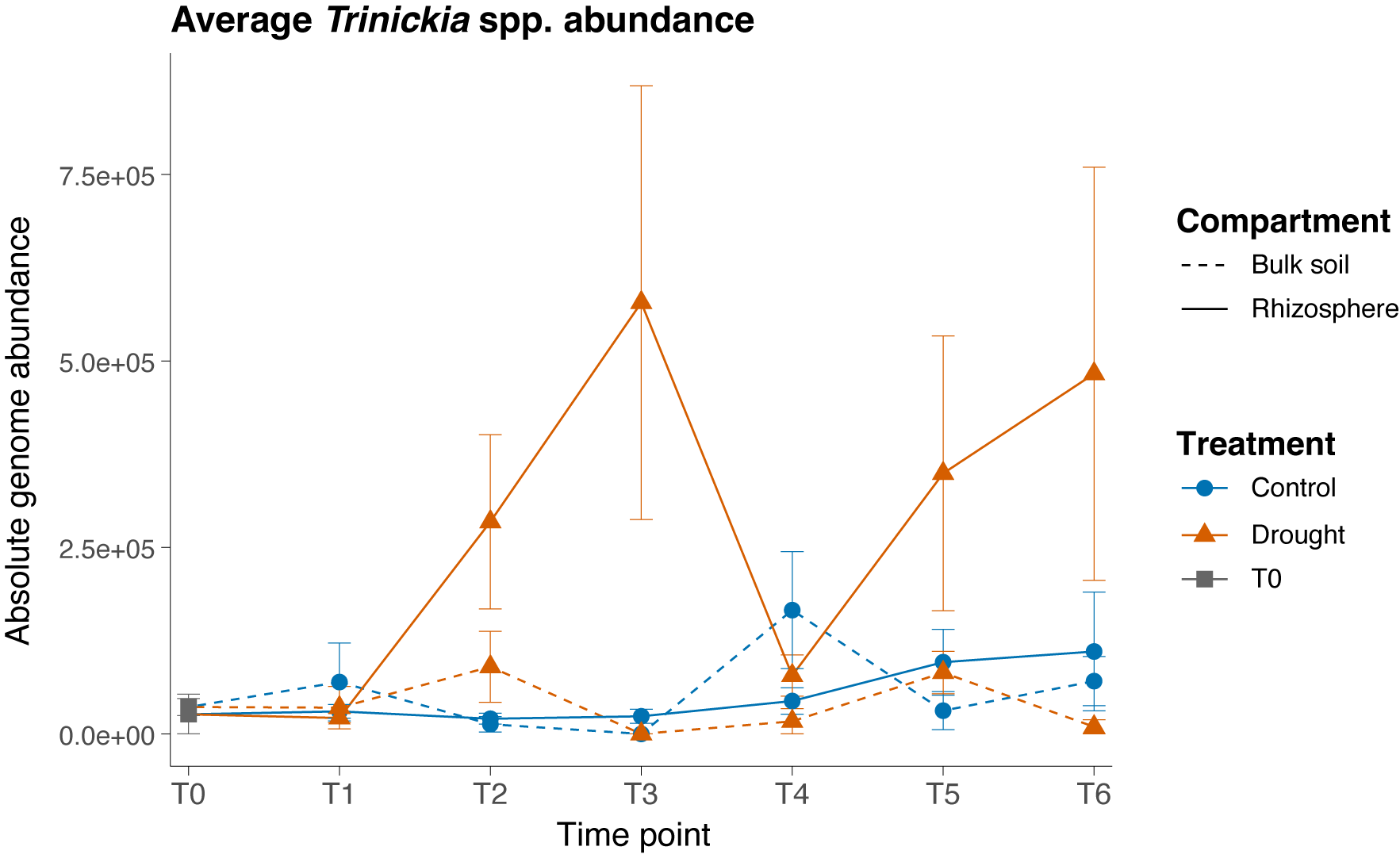

**Fig. S8. Absolute genome abundance of *Trinickia* metagenome-assembled genomes.** Points indicate average absolute genome abundances (± 1 S.E.) for *Trinickia* spp. (*n*=11) at each time point for control (blue circles) and drought-treated (orange triangles) samples, and for rhizosphere (solid lines) and bulk soil (dashed lines) compartments.

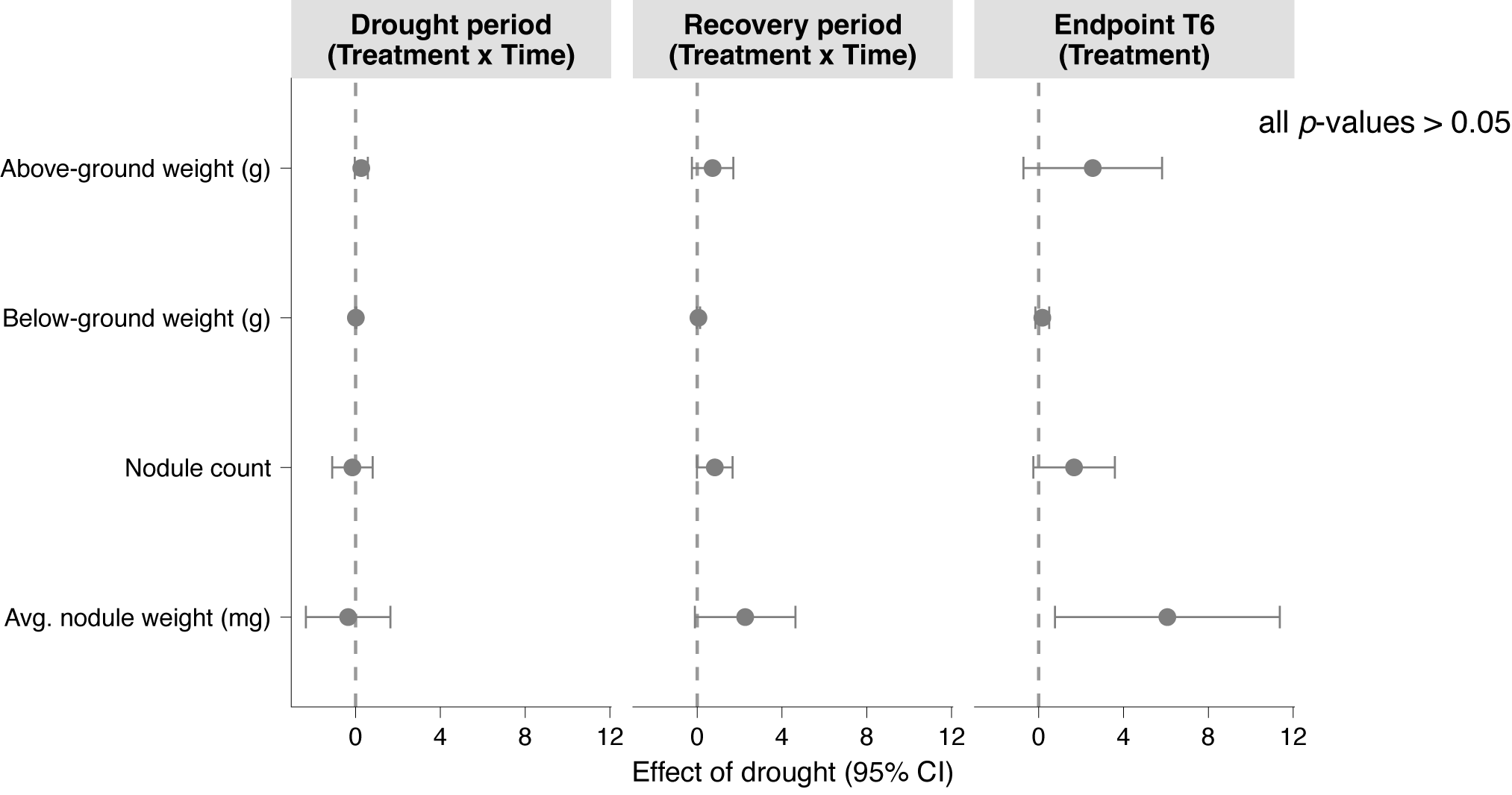

**Fig. S9. Effect of drought treatment on plant traits (n = 6 replicates per treatment per time point).** Drought effects on plant traits were estimated during the drought period (T0 – T3), the recovery period (T3–T6, re-zeroed to a T3 baseline), and at the recovery endpoint (T6). For the drought- and recovery-period models, points represent the Treatment × Time interaction between drought and control plants over that window; for the endpoint model, points represent the direct Treatment effect comparing drought and control plants at T6. Error bars show 95% confidence intervals; the dashed vertical line marks no effect (estimate = 0). No trait showed a statistically significant drought effect (all *p* > 0.05).

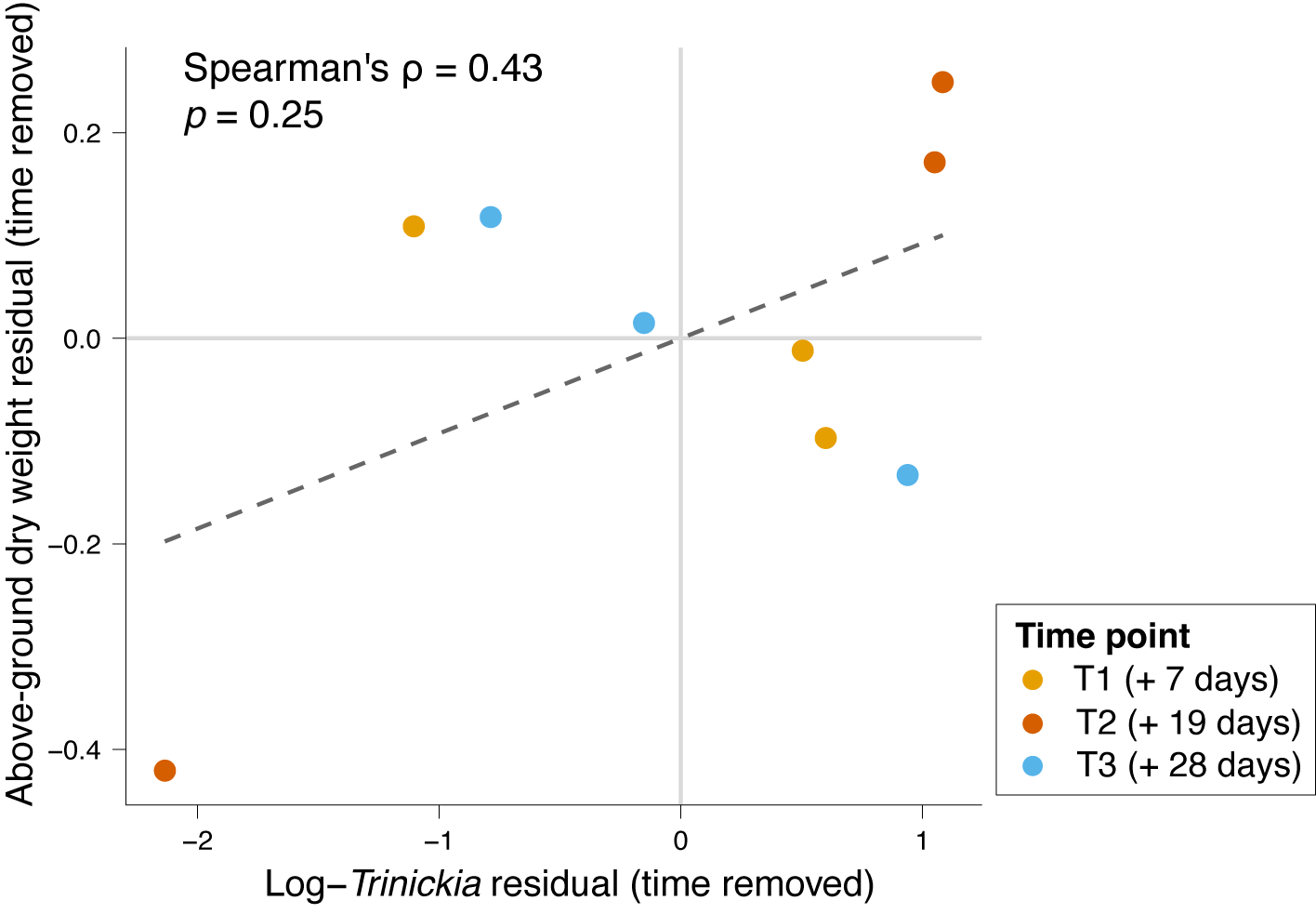

**Fig. S10. *Trinickia* relative abundance and plant host above-ground weight.** Time-adjusted trend between the natural log of *Trinickia* relative abundance and above-ground dry weights, tracked across individual drought sampling time points (T1, T2, and T3). Axes show time-removed residuals to show the underlying relationship between *Trinickia* relative abundance and dry shoot weight once the confounding effects of general plant growth over time is statistically removed (Spearman’s ρ = 0.73, *p* = 0.025; dashed line indicates linear fit).
